# Aryl-hydrocarbon receptor in smooth muscle cells protect against dioxin induced adverse remodeling of atherosclerosis

**DOI:** 10.1101/2024.09.24.614572

**Authors:** Guyu Tracy Qin, Quanyi Zhao, Ayaka Fujita, Isabella Damiani, Meena Easwaran, Sugandha Basu, Wenduo Gu, Daniel Y. Li, Matthew Worssam, Brian Palmisano, Joao Pinho Monteiro, Markus Ramste, Ramendra Kundu, Trieu Nguyen, Chong Park, Chad S. Weldy, Paul Cheng, Juyong Brian Kim

## Abstract

**Introduction:** Environmental exposure to dioxin has been linked to increased myocardial infarction. Smooth muscle cells (SMC) in the coronary vasculature play a critical role in atherosclerotic plaque remodeling due to their phenotypic plasticity, however, the detailed mechanism linking dioxin exposure to adverse SMC modulation is not well understood.

**Methods:** Single-cell RNA and ATAC sequencing and histological analyses were performed on the aorta from mouse models of atherosclerosis exposed to 2,3,7,8-Tetrachlorodibenzo-p-dioxin (TCDD) or control. Primary human coronary artery SMC (HCASMC) treated in culture with TCDD were used to perform RNA-Seq, ATAC-Seq, and functional phenotypic assays. ChIP-Seq was performed with antibodies against Aryl-hydrocarbon receptor (AHR) and TCF21, two of known SMC modulating transcription factors.

**Results:** Modulated SMC were the most transcriptionally responsive cell type to dioxin in the atherosclerotic aorta. Dioxin accelerated disease phenotype by promoting a modulated SMC phenotype early, resulting in increased lesion size, migration of SMC, and macrophage recruitment to the lesion. We found *C3* expressing modulated SMCs to be likely contributing to the increased macrophage recruitment and inflammation. Analysis of the RNA-Seq data from HCASMC treated with TCDD showed differential enrichment of biological pathways related to cell migration, localization, and inflammation. Furthermore, ATAC-Seq data showed a significant activation for pathways regulating vascular development, cell migration, inflammation, and apoptosis. With TCDD treatment, there was also enrichment of AHR ChIP-Seq peaks, while the TCF21 enrichment decreased significantly. The SMC-specific *Ahr* knockout resulted in increased oxidative stress in SMC, increased lesion size and macrophage content, and loss of SMC lineage cells in the lesion cap when exposed to TCDD, consistent with a more vulnerable plaque phenotype.

**Conclusion:** Dioxin adversely remodels atherosclerotic plaque by accelerating the SMC- phenotypic modulation, and increasing inflammation and oxidative stress resulting in increased macrophage recruitment and lesion size. Dioxin may adversely affect the SMC phenotype and disease state by affecting the TCF21 occupancy in the open chromatin regions. Furthermore, we observed that SMC-specific deletion of *Ahr* in mice resulted in worsening of dioxin mediated SMC modulation and atherosclerosis, suggesting that *Ahr* in SMC confers protection against dioxin by promoting a stable plaque phenotype and reducing dioxin induced oxidative stress.

**Summary:**

- Exposure to dioxin, an environmental pollutant present in tobacco smoke and air pollution, accelerates smooth muscle cell modulation, and atherosclerosis.
- Dioxin exposure leads to inflammatory smooth muscle cell phenotype characterized by complement pathway activation and increased macrophage recruitment to plaque
- Aryl-hydrocarbon receptor in SMC protects against oxidative stress, and promotes a stable plaque phenotype

## Introduction

Coronary artery disease (CAD) and its resulting sequelae are the leading cause of morbidity and mortality worldwide.^1^ While genetic and lifestyle factors have been extensively studied in relation to atherosclerosis, emerging evidence highlights the substantial impact of environmental factors.^2, 3^ Among these, exposure to environmental pollutants, such as tobacco smoke, air pollution, heavy metals, and persistent organic pollutants (POPs), has been associated with a heightened risk of atherosclerosis and cardiovascular diseases (CVDs).^4–6^ Tobacco users, in particular, face a risk more than double that of non-users, with smoking being attributed to one in four global cardiovascular disease-related deaths.^7^ Furthermore, air pollution has been established as a significant factor in increasing the risk of CAD.^8^ Despite mounting evidence, the intricate molecular mechanisms through which environmental toxicants induce and exacerbate atherosclerosis have yet to be fully elucidated. Complicating matters further, the interplay between genetics and the environment (GxE) substantially impacts disease susceptibility.^9, 10^ Nonetheless, studying this interaction has been challenging due to limited statistical power and a lack of well-documented individual-level measurements of exposures.^11, 12^

In this context, dioxins and related compounds have emerged as well-characterized environmental toxins, constituting a significant part of cigarette smoke and air pollution.^13–15^ These compounds, including 2,3,7,8-tetrachlorodibenzo-p-dioxin (TCDD), acts by activating the AHR transcription factor and its downstream genes but also by non-canonical pathways to turn on detoxification program as wells as mediate immune function and cell proliferation.^16, 17^ The AHR, upon ligand exposure, triggers downstream gene transcription program, including activation of the cytochrome p450 enzymes. Epidemiological studies have consistently associated environmental dioxin exposure with cardiovascular diseases such as atherosclerosis and myocardial infarction.^18, 19^ For example, the Seveso Women’s Health Study (SWHS), which followed women exposed to TCDD during an industrial accident in Seveso, Italy, found a significant increase in ischemic heart disease among the exposed population.^20^ These large longitudinal studies collectively highlight the significant public health impact of dioxin exposure on cardiovascular disease development. However, the specific molecular mechanism by which dioxin may induce increased atherosclerosis in the vascular wall remains unknown.

To address this knowledge gap, this paper aimed to explore the molecular mechanisms through which dioxin contributes to disease progression in atherosclerosis. Smooth muscle cells (SMC), a major component of the vascular wall, plays a crucial role in disease pathogenesis through epigenetic control of phenotypic plasticity, including unstable plaque architecture implicated in plaque rupture events leading to myocardial infarction and stroke.^21, 22^ Genome-wide association studies have also identified numerous genes that likely modify disease risk through SMCs.^23–25^ Previously, our research demonstrated that the *AHR* gene can modulate SMC function and influence atherosclerosis development in mouse models of the disease.^26, 27^ Building upon this foundation, the current study sought to provide valuable insights into the intricate relationship between environmental dioxin exposure, SMCs, and atherosclerosis. Using detailed characterization of the dioxin-induced atherosclerotic lesion employing single-cell sequencing and histology, we reveal that dioxin exposure accelerates the SMC phenotypic modulation increasing SMC participation in the atherosclerotic lesion, and promotes an inflammatory milieu increasing macrophage recruitment to the lesion. Furthermore, we explored the role of *AHR* gene in how SMC responds to dioxin exposure, and conclude that *AHR* in SMC confers protection against dioxin induced atherosclerosis by maintaining a stable SMC-rich lesion cap and decreasing burden of oxidative stress.

## Methods

All data and materials are deposited to the National Center for Biotechnology Information Gene Expression Omnibus (GSE276145, GSE276150, GSE276152, GSE276870 and GSE276872). All methods and materials used in this study are described in detail in the Materials and Methods section of the Data Supplement. In the following sections, please see a brief summary of the most relevant details.

### In vitro genomic and functional studies in primary and immortalized human coronary artery smooth muscle cell (HCASMC)

Primary HCASMC were purchased from Cell Applications, Inc, and experiments were performed with HCASMCs between passages 5 and 8. Immortalized HCASMC was generated using introduction of hTert vector (Addgene #85140). A full description of the methodology for the culture of human coronary artery SMCs (HCASMCs), and in vitro genomic and functional studies is provided in the Supplemental Methods.

### Mouse models

The animal study protocol was approved by the Administrative Panel on Laboratory Animal Care at Stanford University. The DRE-LacZ strain was rederived from cryopreserved embryos at the Jackson Laboratories (B6.Cg-Tg(DRE-lacZ)2Gswz/J strain, JAX #006229)^28^ then bred to ApoE–/– background (B6.129P2-*Apoe^tm1Unc^*/J, JAX #002052). SMC-specific lineage tracing and *Ahr* gene knockout in the atherosclerotic model was performed as described previously.^27^ Tamoxifen was administered by oral gavage at 8 weeks of age to induce the lineage marker and *Ahr* knockout, followed by the initiation of a high-fat diet (Dyets No. 101511).

### Murine aortic tissue processing and histology

Immediately after sacrifice, mice were perfused with 0.4% paraformaldehyde and aortic tissue was harvested, embedded in optimal cutting temperature compound, and sectioned. Immunofluorescence and immunohistochemistry were performed and lesion areas profiled as described previously.^27, 29^ A full description of staining and quantification methods is provided in the Supplemental Methods.

### In vitro and In vivo exposure

TCDD was introduced to culture for final concentration of 10nM for all *in vitro* experiments including RNA-seq, ATAC-seq, ChIP-Seq, and SMC phenotypic assays. CSE was introduced at 10% v/v into the media for treatment of HCASMC. For in vivo exposure, a single intraperitoneal injection of 15μg/kg TCDD was administered at initiation, followed by a weekly maintenance dose of 1μg/kg TCDD as described previously.^30^ Control mice received saline injections.

### Analysis of single-cell sequencing data

A full description of the methodology for the acquisition and analysis of both single-cell RNA sequencing (scRNA-Seq) and single-cell assay of transposase-accessible chromatin with sequencing (scATAC-Seq) data is provided in the Supplemental Methods.

### Statistical methods

Differentially expressed genes in the scRNA-seq and scATAC-seq data were identified using a Wilcoxon rank-sum test. For overlapping of genomic regions or gene sets, we used the Fisher exact test to test for enrichment. For comparison of cell cluster proportions, Chi-square test was used. For comparisons between 2 groups of equal variances, an unpaired 2-tailed *t* test was performed. For multiple comparisons testing, 1-way ANOVA accompanied by a Tukey post hoc test were used as appropriate. All error bars represent standard error of the mean. Number of asterisks for the P values in the graphs: ****P<0.0001, ***P<0.001, **P<0.01, *P<0.05.

## Results

### Modulated smooth muscle cells have the strongest dioxin response activity in the vascular wall

Utilizing dioxin response element LacZ reporter transgenic mouse model of atherosclerosis (DRE-LacZ Tg/ApoE KO), we have previously found robust dioxin response element (DRE) activity in the vascular wall (**Figure S1A**).^27^ To assess the effect of dioxin on the vascular wall at a single-cell resolution, we performed a single-cell RNA-seq of the atherosclerotic aortic sinus of the DRE-lacZ Tg/ApoE KO mice on HFD for 10 weeks and subsequently exposed to vehicle (Control, n=3) or short term dioxin (TCDD, n=3) for 2 weeks with HFD.

Analysis of the scRNA-seq identified previously described cell types including SMC and modulated SMC (Mod-SMC) populations based on PCA based clustering (**Figure 1A; Figure S1B**). We looked at the canonical downstream gene of DRE activation by Ahr, *Cyp1b1*, as a marker of overall response to TCDD. To identify the most dioxin responsive cell type within the vasculature, we compared the weighted fold change (FC) of *Cyp1b1* expression (i.e. FC (Average Expression) x FC (% cells expressing gene in cluster)) between different cell types of the vascular wall (**Figure 1B**). The Mod-SMC population stood out as the most responsive cell type based on *Cyp1b1* expression (6.5 fold increase in average expression; 4.5 fold increase in % expressing cells), followed by SMC and fibroblast populations. A closer look at the dioxin-induced transcriptional changes in the Mod-SMC using differential expression analysis of control and dioxin exposed group identified several key pathways implicated in atherosclerosis including TGF-β signaling, VEGF production, cell migration and proliferation, inflammatory and oxidative response, and ECM and collagen fibril organization pathways (**Figure 1C, Table S1**).

**Figure 1.**
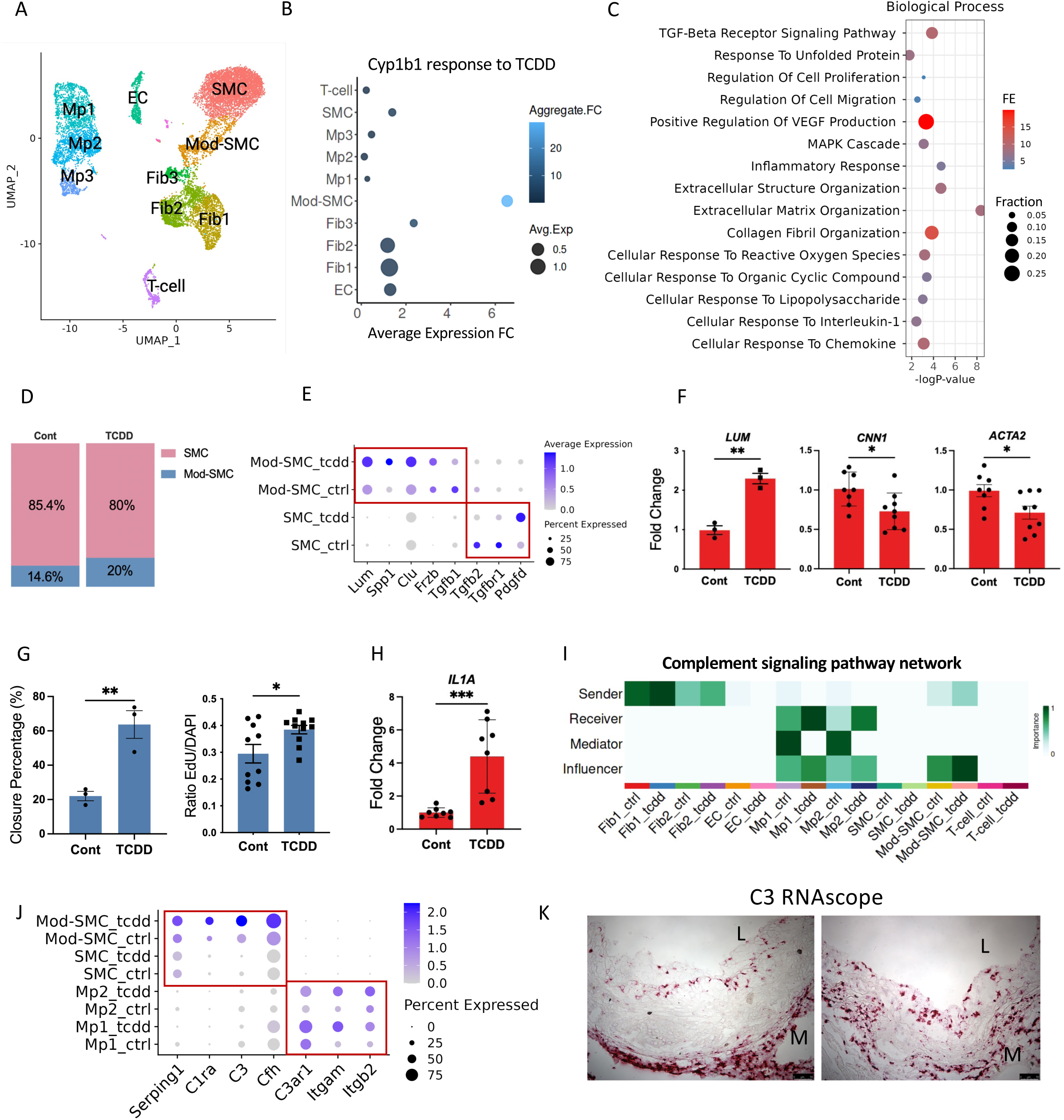
Dioxin induces significant transcriptional and phenotypic response in the modulated smooth muscle cells present in atherosclerosis. A. UMAP of scRNA-seq from aorta of DRE-lacZ Tg; ApoE KO mice on 12 weeks of HFD, exposed to 2 weeks of TCDD (SMC = smooth muscle cells; Mod-SMC = Modulated SMC; Fib = Fibroblast; EC = Endothelial cells; Mp = Macrophage). B. Modulated SMC displayed the greatest fold change in average expression of *Cyp1b1*, and aggregate response to TCDD (product of average expression fold change and fold change in % cells expressing *Cyp1b1*). C. Gene ontology pathway analysis of differentially expressed genes in Mod-SMC identify significant alterations in inflammatory/MAPK/IL-1, collagen/ECM organization, and TGF-beta receptor pathways. FE = Fold Enrichment; Fraction = # of DE genes present within the ontology defining gene set/# of Total DE genes. D. Dioxin exposure shift larger proportion of the SMC lineage cells toward the modulated SMC transcriptional phenotype in the atherosclerotic lesion (Chi-square p < 2.2e-16). E. Transcriptional markers of modulated SMC are upregulated in response to TCDD treatment in Mod-SMC (*Lum, Spp1, Clu, Frzb*), and *Pdgfd*, a CAD gwas gene we have previously found to regulate SMC migration and proliferation, is upregulated in SMC. Tgf-β pathway genes are downregulated in SMC and Mod-SMC. F. Treatment of HCASMC with TCDD showed increase in FMC marker *LUM* (p=0.0016), and downregulation of SMC markers including *CNN1* and *ACTA2* (p=0.021, p=0.027, respectively). G. TCDD (10nM) induced increased migration (*left*, p=0.008) and proliferation (*right,* p=0.024) of HCASMC *in vitro* measured by EdU incorporation; H. TCDD induces upregulation of IL1A in HCASMC (p=0.0007). I. Cell-cell interaction analysis with *CellChat* found COMPLEMENT pathway showing significant outgoing influence from Mod-SMC, and increased net incoming signaling in macrophages following TCDD exposure. J. Transcriptional markers of complement pathway are upregulated in response to TCDD treatment in Mod-SMC (*Serping1, C1ra, C3, Cfh*), and the complement receptors (*Itgam, Itgb2, C3ar1*) are upregulated in the macrophages. K. RNAscope of *C3* RNA expression in atherosclerotic mouse aortic sinus show expression in the media and the lesion area. There is robust signal in media and intima and the signal is localized to the medial aspect of the lesion cap in the mature lesion (L= Lumen, M=Media).

### Short-term dioxin exposure promotes SMC phenotypic modulation

Comparing the cell cluster proportions following short-term exposure to dioxin (2 weeks), we found a significant transcriptional shift of the SMC-lineage cells towards the Mod-SMC phenotype (**Figure 1D**). Consistent with the increase in proportion of Mod-SMC cells based on PCA, markers of modulated SMC, including *Lum and Spp1* were significantly upregulated in the dioxin-treated group compared to control (**Figure 1E**). Furthermore, several cellular programs known to promote SMC de-differentiation were activated in the dioxin treated group compared to controls. For example, *Pdgfd*, a well-known inducer of SMC de-differentiation was up-regulated in the SMC, while TGF-β signaling, known for its role in SMC phenotypic modulation and ECM regulation in atherosclerosis was down-regulated in SMC and Mod-SMC (*Tgfb1, Tgfb2, Tgfbr)*.

We validated the effect of dioxin on SMC phenotypic modulation *in vitro* using human coronary artery smooth muscle cells (HCASMC). HCASMC treated with 10nM of dioxin for 24 hours resulted in increase in *LUM,* a marker of SMC de-differentiation and decrease in *CNN1*, and *ACTA2*, both markers of quiescent SMC (**Figure 1F**). Dioxin exposure also resulted in changes to cellular phenotypes of HCASMC, including increase in migration and proliferation (**Figure 1G**).

### Dioxin increase pro-inflammatory signal of Mod-SMC leading to increased cross-talk with macrophages

Following exposure to dioxin, the Mod-SMC exhibited increased levels pro-inflammatory gene expression, including *Cxcl12, Lgals3, and Gas6* (**Figure S1C**). HCASMC treated with TCDD *in vitro* also showed increase in marker of inflammation, including *IL1A* (**Figure 1H**). Furthermore, quantitative ROS assay found increase in cellular oxidative stress following dioxin exposure in HCASMC (**Figure S1D**). A *CellChat* analysis was performed to identify potential cell-cell communications with outgoing inflammatory signals from the SMC population that increased with TCDD treatment. Complement signaling pathway was identified as having a clear outgoing signaling from modulated SMC and fibroblasts interacting with the macrophage populations (**Figure 1I**). Closer look at the genes responsible for this interaction found complement pathway genes, including *C1ra, C3, Cfh, Serping1* increase in the modulated SMC following dioxin exposure, while the corresponding complement receptors *C3ar1*, *Itgam*, and *Itgb2* expression increase in the macrophage populations (**Figure 1J**). Spp1 signaling pathway was also strongly induced following dioxin treatment, with Mod-SMC as the key sender cell type towards macrophage and SMC (**Figure S1E**). The expression of *C3* gene was found to be largely restricted to the modulated SMC and fibroblasts, and complement receptors expression restricted to macrophages based on the scRNA-seq (**Figure S1F**). RNAscope of *C3* RNA in the mouse atherosclerotic lesion showed localization of RNA expression to the media as well as the lesion cap (**Figure 1K**).

### Cigarette smoke extract activates the dioxin response pathway in HCASMC resulting in shared gene expression changes with dioxin

To further validate mouse in vivo findings, HCASMC were treated with vehicle and TCDD (10nM) for 24 hours, and resulting RNA was sequenced for transcriptomic profiling. Dioxin treatment generated a significant transcriptional response with 493 genes upregulated, and 412 genes downregulated (FDR <0.05, fold change >1.3) (**Figure S2A; Table S2**). These differentially expressed genes enriched for biological pathways including blood brain barrier transport, regulation of cell proliferation, migration, and angiogenesis, monocyte differentiation, and response to inflammation and hypoxia (**Figure 2A**).

**Figure 2.**
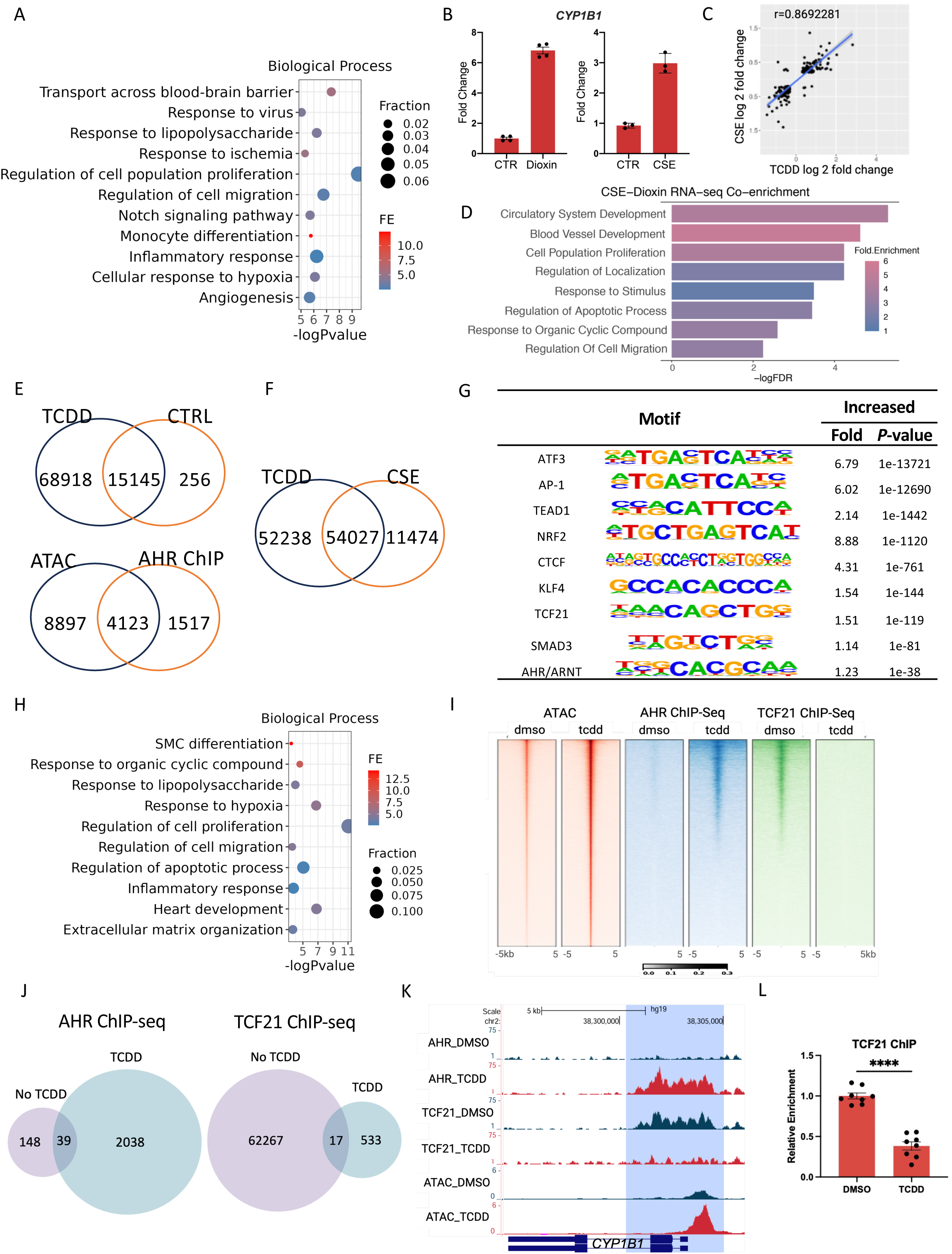
Dioxin induces significant transcriptomic and epigenetic alternations in HCASMC. A. Dioxin induced significant change in the transcriptome of HCASMC resulting in enrichment for GO Biological Pathways including cell communication, migration, proliferation, and differentiation. B. Dioxin and cigarette smoke extracts (CSE) both robustly activate the dioxin response pathway (p<0.0001; p=0.0004, respectively); C. Comparison of the RNA-seq analysis of HCASMC treated with TCDD and CSE show significant correlation (r=0.87). D. Genes that are differentially regulated with both CSE and dioxin enriched for pathways of circulatory system development, cell proliferation, regulation of localization/migration. E. (*top*) TCDD resulted in significant increase in the chromatin accessibility landscape of HCASMC resulting in 68918 differentially accessible peaks compared to only 256 peaks in control (LR >3, FDR<0.05). (*bottom*) There was also significant overlap of genes proximal to the open chromatin region and AHR binding sites from previously performed ChIP-seq (Fisher’s test, p<e-255). F. There is significant overlap of chromatin accessible peaks between TCDD and CSE treatment in HCASMC (Fisher’s, p<e-255). G. The TCDD responsive open chromatin regions enriched for key SMC TF motifs including KLF4, TCF21, SRF, SMAD3, and AHR/ARNT H. The genes common to the open chromatin regions from TCDD treatment and the AHR ChIP-seq peaks enriched for pathways regulating cell proliferation and migration, extracellular matrix organization, differentiation, and response to organic cyclic compound, hypoxia and inflammation. I. There is significant increase in chromatin accessibility following dioxin treatment (red). Centered around the ATAC-seq peaks, there is increased AHR binding (blue), and a reciprocal genome-wide reduction of TCF21 binding at regions following TCDD treatment (green). J. A significant drop in TCF21 binding is observed following TCDD treatment. *macs bdgdiff* identified 2038 AHR differential binding peaks when treated with dioxin, while only 148 peaks without dioxin (*left*); for TCF21 ChIP-Seq there were 62267 TCF21 binding peaks in the absence of TCDD compared to only 533 peaks when treated with TCDD (*right*). K. In the promoter region of *CYP1B1*, chromatin accessibility and AHR binding increased, while TCF21 binding decreased following TCDD treatment. L. ChIP-qPCR of the region confirmed significant reduction in TCF21 binding following TCDD treatment.

In order to compare the effect of cigarette smoke with the dioxin response, we treated HCASMC with aqueous cigarette smoke extract (CSE) generated using the Scireq inExpose system (Montreal, QC, Canada). Exposure of HCASMC to CSE resulted in robust activation of the dioxin responsive gene *CYP1B1* (**Figure 2B**). We subsequently conducted RNA-seq of HCASMC exposed to CSE to compare the transcriptional response with dioxin. There were significant transcriptional changes following treatment with CSE (316 upregulated and 220 downregulated genes; Fold change >1,3, FDR <0.05; **Figure S2A; Table S3**). Upstream transcription factors predicted to regulate these changes included key SMC TFs (eg. KLF4, SMAD3, AHR) as well as inflammatory TFs, including STAT2/3 and NFKB1/RELA (**Figure S2B**). Differentially regulated biological pathways included ER stress, circulatory system development, cell localization, ECM, and response to cytokine stimulus and oxidative stress (**Figure S2C**). We found highly significant correlation (R = 0.87, **Figure 2C**) and a significant overlap in the differentially regulated gene list between CSE and dioxin (**Figure S2D**) suggesting that a significant portion of CSE effect in HCASMC is driven by the dioxin response pathway. These overlapped genes enriched for pathways regulating circulatory system development, cell migration and proliferation, and response to stimulus (**Figure 2D**).

### Dioxin increases chromatin accessibility in HCASMC and regulates its epigenetic state

We further investigated the role of dioxin in regulating the epigenetic landscape of HCASMC. ATAC-seq was performed to assess the changes in chromatin accessibility following dioxin exposure. TCDD acted as an overall activator resulting in 68918 upregulated peaks compared to control condition (*macs2 bdgdiff*, FDR<0.05, log10(likelihood ratio) >3, **Figure 2E (**top)).

These upregulated peaks were in proximity of 8897 genes, which significantly overlapped with genes (n=4123) proximal to AHR binding sites identified on AHR ChIP-Seq performed with TCDD treated HCASMC (**Figure 2E** (bottom)). The upregulated peaks enriched for pathways including cellular response to stress, vasculature development, response to cytokine stimulus, and ECM organization (**Figure S2E**). We also performed an ATAC-seq of HCASMC treated with CSE. Overall, CSE exposure increased chromatin accessibility in HCASMC at shared genomic locations with TCDD exposure, and differential accessibility (DA) peaks enriched for pathways including vasculature development, regulation of cell migration, angiogenesis, MAPK/NF-kB signaling, ER stress and apoptosis (**Figure S2G, S2H**). When comparing the DA peaks with TCDD treatment, there was a significant overlap between TCDD and CSE treatment (**Figure 2F, Figure S2F**).

Furthermore, we identified the overrepresented motifs in the differentially enriched chromatin peaks in response to TCDD. This analysis found motifs of key SMC TFs including AP-1, TCF21, AHR/ARNT, KLF4, SMAD3 and SRF, as well as ATF3, NRF2/NFE2, and TEAD enriching in the differentially accessibly (DA) chromatin regions (**Figure 2G**). The genes in the regions of DA chromatin peaks that are also differentially expressed following TCDD exposure enriched for pathways involved in vasculogenesis, response to inflammation and organic cyclic compound, cell proliferation and migration, cell differentiation (**Figure 2H**).

### Dioxin results in global reduction of TCF21 binding to chromatin

We have previously shown that TCF21 interacts with AHR to activate an inflammatory gene expression program that is exacerbated by environmental stimuli.^26^ We have also found that AHR and TCF21 binding loci co-localize genome-wide. To understand the interaction of TCF21 and AHR in response to dioxin, we performed AHR and TCF21 ChIPseq treated with or without TCDD. As expected, we found a significant increase in total AHR ChIP peaks when treated with TCDD compared to control (2038 and 148, respectively), and genome-wide colocalization of the peaks with TCDD ATAC-seq increased peaks (**Figure 2I, 2J** (*left*), **S2I**). However, we found a significant genome-wide reduction in the chromatin occupancy of TCF21 following TCDD treatment (62267 peaks without TCDD and 533 with TCDD respectively, **Figure 2J (***right***), S2I**). Evaluation of the *CYP1B1* region showed increased chromatin accessibility, increased AHR binding, and decreased TCF21 binding at the promoter region following TCDD exposure (**Figure 2K**). To confirm our findings, we performed ChIP-qPCR using TCF21 antibody at the *CYP1B1* locus, observing a significant reduction in TCF21 binding following TCDD treatment (**Figure 2L**). These findings demonstrate that dioxin exerts a significant epigenetic effect on SMC by modulating chromatin accessibility and key SMC transcription factor-DNA interaction.

### Prolonged dioxin exposure leads to increased cellular stress and reduced proportion of modulated SMC in mice

To further characterize the modulation of the SMC state following chronic exposure to dioxin, we employed single-cell RNA sequencing (scRNA-seq) using SMC-lineage tracing (LnT) models as described previously.^27^ The mice (n=6 in each group) were exposed to HFD only for 16 weeks (WT), or 8 weeks of HFD followed by TCDD and HFD for 8 weeks (TCDD (8wk)) (**Figure S3A**). Samples from 3 mice were merged for each library preparation. The tdTomato+ and tdTomato-population of cells were separated by FACS and 10x single-cell library were generated for sequencing.

After QC, we identified 4 main clusters within the tdTomato+ population of cells (n= 11,234), which were identified as SMC1, SMC2, fibroblast-like phenotype fibromyocyte (FMC) and chondrogenic phenotype chondromyocyte (CMC) (**Figure 3A**). Genes that distinguished these clusters included mature SMC markers such as *Tagln*, *Acta2* and *Cnn1*, fibroblast-like marker Lum, and chrondrogenic marker *Spp1* and *Col2a1* (**Figure S3B**). TCDD-induced *Cyp1b1* gene was highly expressed in FMC compared with WT (**Figure S3C**). Overall, we found that TCDD induced a significantly larger proportion of SMCs and reduced proportions of modulated SMC including FMC and CMC (**Figure 3B**).

**Figure 3.**
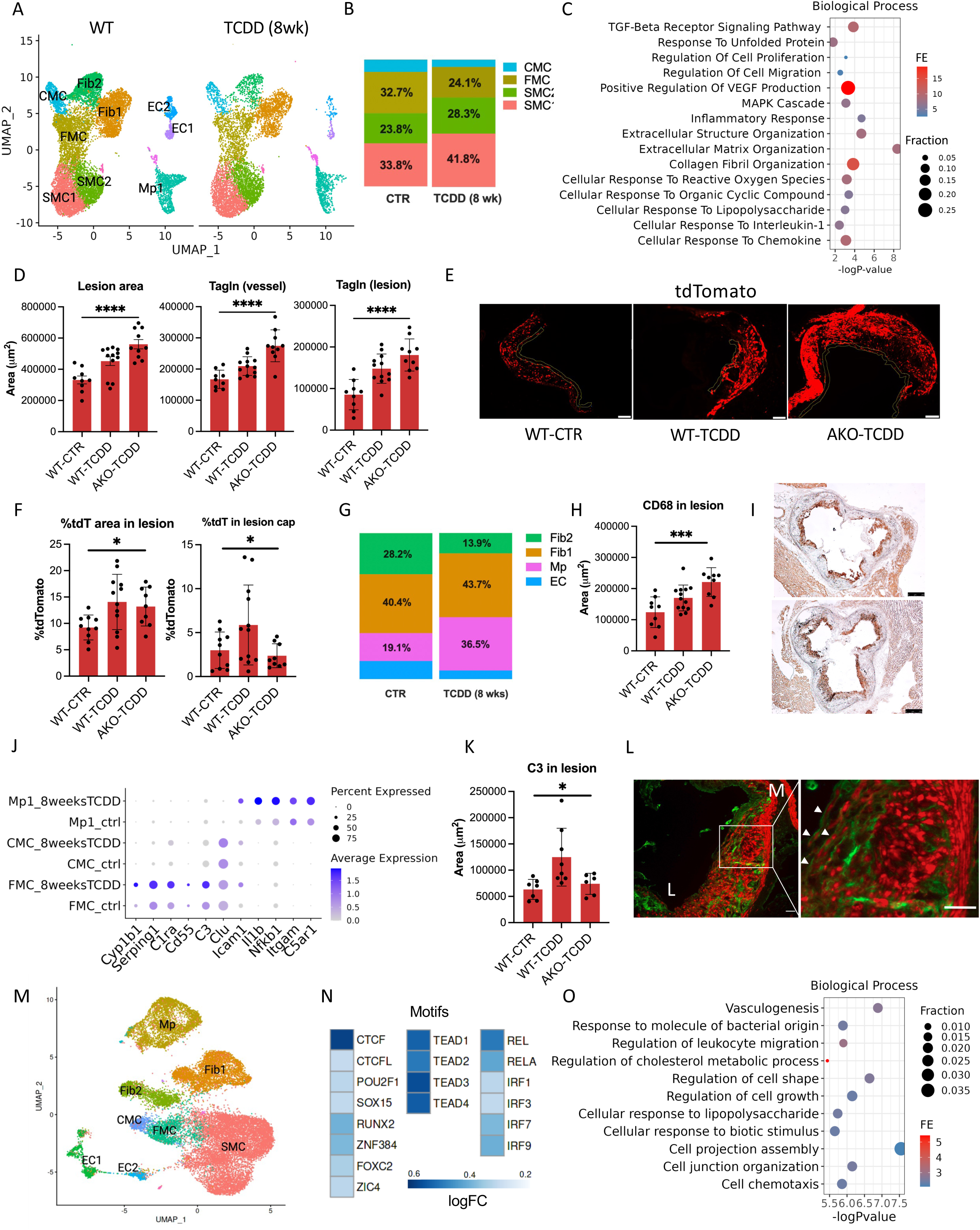
Dioxin induce SMC phenotypic modulation and increased inflammatory burden. (A) UMAP projection of scRNA-seq PCA clustering from Myh11-Cre, Rosa26-tdT, ApoE null mice treated with 16 weeks of HFD (CTR, n=3) or 8 weeks HFD followed by 8 weeks of HFD and TCDD (TCDD, n=3) from 8 weeks of age (**Figure S3A**). (B) There is significant shift in the proportions of tdTomato-positive population of cells, including increase of SMC1 (33.8% to 41.8%) and SMC2 (23.8% to 28.3%) clusters, and decrease in FMC (32.7% to 24.1%) and CMC (9.7% to 5.6%) clusters (Chi-square p < 2.2e-16). (C) The differentially regulated genes in FMC cluster in response to TCDD enrich for cell proliferation/migration/adhesion, chondrocyte differentiation, ECM organization and ER stress. (D) A histological analysis of the aortic root atheroma was performed on *SMC-LnT* mice treated with 16 weeks of HFD (WT-CTR), 16 weeks of HFD and TCDD (WT-TCDD), and SMC-AhrKO mice treated with 16 weeks of HFD and TCDD (AKO-TCDD). (*left*) There was overall increase in the lesion area in the WT-TCDD group compared to control. The total lesion area was greater in the AKO-TCDD group compared to WT-TCDD (ANOVA p<0.0001). (center) There was overall increase in Tagln+ SMC in the vessel and (right) lesion area compared to WT-CTR. There was a trend toward further increase in *Tagln+* area in the lesion of AKO-TCDD group compared to WT-TCDD (ANOVA p<0.0001). (E) The tdTomato migration and localization was quantified in the atherosclerotic lesions (scale bar =50um). (F). (*left*) There was increase in tdT+ area in the lesion of WT-TCDD and AKO-TCDD compared to WT-CTR (*ANOVA p=0.025*), and (*right*) increase in the tdT signal in the lesion cap in the WT-TCDD compared to WT-CTR. A significant reduction of tdT+ area was observed in lesion cap of the AKO-TCDD group compared to WT-TCDD (ANOVA, *p=0.035*). (G) The proportion of macrophages within the non-tdTomato cells from the scRNA-seq data show significant increase in the macrophage population following TCDD exposure (Chi-square p < 2.2e-16). (H) There is significant increase in the Cd68+ area in lesion by IHC in the TCDD group compared to CTR, and further increase in the TCDD+AKO group (ANOVA p=0.0004). (I) Aortic sinus with Cd68 immunohistochemistry (J) The complement associated pathway genes including *C1ra*, *C3*, and *Serping1*, as well as complement receptors and other inflammatory markers genes including *Il1b, Nfkb1, and Icam1* are upregulated following TCDD exposure. (K) There was increase in the total C3 area in the lesion following TCDD exposure (WT-CTR vs. WT-TCDD, p=0.014), however, no increase was seen in the AKO-TCDD group (ANOVA p=0.012). (L) Immunofluorescence of C3 (green) show significant expression of C3 protein in the atherosclerotic lesion, and colocalize at the cell surface of cells expressing tdTomato (red) signal in the lesion cap, as well as intima of lesion. (right) closer look shows C3 expression on the cell surface of the tdTomato+ SMC-lineage cells at the lesion cap (arrowheads). (M) UMAP projection of the scATAC-seq of SMC-LnT mice exposed to 16 weeks of HFD or HFD+TCDD with clusters identified via label transfer from scRNA-seq. The SMC1 and SMC2 clusters and MP1 and MP2 clusters were merged to SMC and MP clusters, respectively. (N) Motifs including IRF1/3/7/9, TEAD, CTCF, and RELA are significantly enriched in the FMC population after TCDD treatment. (O) Enrichment analysis of the dioxin responsive chromatin accessibility peaks identify pathways including vasculogenesis, cell chemotaxis and growth, and leukocyte migration.

TCDD induced 469 upregulated DE genes and 43 downregulated DE genes in the FMC cell cluster (p<0.05, log2FC>0.25, **Figure S3D; Table S4**). These genes enriched for TGF-β signaling pathway, regulation of cell migration and proliferation, regulation of VEGF production, inflammatory response and MAP cascade, response to chemokine/IL-1, and reactive oxygen species, extracellular matrix organization, collagen fibril organization, endoplasmic reticulum stress and unfolded protein response (**Figure 3C**). Significant changes in gene expression levels were also seen in the SMC and MP clusters (**Table S4**).

### Chronic exposure to dioxin increases atherosclerosis and affects the SMC participation in lesion remodeling

In order to assess the impact of chronic dioxin exposure on atherosclerosis formation and smooth muscle cell remodeling, we employed the SMC-lineage tracing mouse model on hyperlipidemic background (Myh11^CreERT2^, Rosa^tdT/tdT^, ApoE-/-; SMC-LnT) for histological analysis of lesion characteristics. We compared the SMC-LnT mice exposed to TCDD and HFD for 16 weeks (WT-TCDD, n=13) with the vehicle treated SMC-LnT mouse group given HFD only (WT-CTR, n=10). In order to assess the role of *Ahr* in mediating SMC response to dioxin, we also extended the histological study of the lesion characteristics to the previously described SMC-lineage specific *Ahr* deletion model (Myh11^CreERT2^, Ahr^ΔSMC/ΔSMC^, Rosa^tdT/tdT^, ApoE-/-; SMC-AhRKO) exposed to 16 weeks of TCDD and HFD (AKO-TCDD, n=10).

We performed immunohistology of aortic root and characterized the atherosclerosis lesions for WT-CTR, and WT-TCDD. No significant differences in weight or plasma lipid levels were identified between groups. We found TCDD increased the total lesion area compared with CTR (**Figure 3D).** The Tagln staining revealed that the overall SMC areas in both the vessel and lesion were significantly increased with TCDD injection compared with WT (**Figure 3D**). We also quantified the tdTomao+ signal in the diseased vessel (**Figure 3E**). After TCDD exposure, there was significant increase in the proportion of tdTomato+ cells in the lesion and lesion cap (**Figure 3F**). These findings suggest that dioxin exposure increases SMC proliferation and migration to lesion and lesion cap. In the SMC-AhRKO treated with 16 weeks of HFD/TCDD (AKO-TCDD), there was further increase in the total lesion area and Tagln+ area in lesion compared to WT-TCDD (**Figure 3D, S3F**). Despite increase in Tagln+ plaque area in AKO-TCDD, we found a significant decrease in the proportion of tdTomato+ cells in the lesion cap area of AKO-TCDD group compared to the WT-TCDD group (**Figure 3F**).

### Dioxin increases inflammatory burden of atherosclerosis and epigenetic changes in SMC

Analysis of the non-tdTomato portion of the digested aortic sinus showed significant increase in the macrophage proportion following TCDD treatment (**Figure 3G**). We found an increase in the Cd68 positive area in the lesion of the TCDD treated group compared to the control group, consistent with the findings from the single-cell analysis (**Figure 3H, I**). There was further increase in the Cd68 area of the Ahr-KO group compared to the TCDD group.

Upon examination of scRNA-seq data, we observed increased expression of complement pathway genes, including *C3* gene in the FMC population along with complement receptor genes (*Itgam, C5ar1*) and proinflammatory genes (eg. *Il1b*, *Nfkb1*, *Icam1*) in the macrophage population after chronic TCDD injection (**Figure 3J**). Consistent with these findings, on histology the C3-positive area and C11b(Itgam)-positive areas was significantly increased in size following TCDD exposure and there was a trend towards increase in C11b(Itgam)-positive area (**Figure 3K, Figure S3G**). Closer look at the C3 protein expression show C3 localized at the surface of the tdTomato+ cells of the lesion cap, consistent with Mod-SMC expression of C3 in the atherosclerotic plaque (**Figure 3L**). Immunohistochemistry of the C3 and Itgam showed adjacent localization of the tdT+/C3+ cells and the Itgam+ cells in the atherosclerotic plaque potentially indicating that Itgam+ macrophages are attracted to the C3+ Mod-SMC. (**Figure S3I**)

We then performed a single-cell ATAC-seq (scATAC-seq) of the atherosclerotic aorta from mice treated with 16 weeks of HFD and 16 weeks of HFD/TCDD (WT-CTR and WT-TCDD, respectively). Chromatin accessibility at the Cyp1b1 promoter region increased as expected (data not shown). To help interpret the scATAC-seq data, we classified cells based on label tranfer from scRNA-seq experiment clustering (**Figure 3M)**. There were significant differences in cell clustering between the treatment conditions (**Figure S4A**).

The comparative analysis of scATAC-seq from the WT-CTR and WT-TCDD group found significant changes in the chromatin accessibility signatures following dioxin treatment. In the FMC, there were 1186 peaks with increased accessibility and 586 with decreased accessibility following TCDD exposure (p<0.05) (**Table S5**). The activated chromatin regions in the FMC cell population enriched for transcription factor motifs including IRF1/9/7/3, CTCF, and FOXC2 (**Figure 3N**). The pathway analysis based on proximity to genes found enrichment for pathways including vasculogenesis, regulation of leukocyte migration, cell chemotaxis, adhesion, and cell growth (**Figure 3O**). Similarly, the activated regions in the SMC cluster of cells enriched for CTCF/CTCFL, MEF2A/B/C, TEAD, IRF1, NRF1, and YY2, while the activated regions in macrophages enriched for ETV, ELF, IRF1/7/8/9, CTCF, and JUN (**Figure S3J**). CTCF is a pioneer TF that is known to be a key regulator of cell identify gene transcription with critical function in cardiovascular development.^31, 32^ Increased CTCF binding has been linked to increased proliferation of SMC,^23^ and regulation of SMC phenotype through its effect on SYNPO2 in response to stress.^33^ Interferon response and its downstream effector STAT1:STAT2 are known upstream regulators of innate immune function and inflammation. Furthermore, changes in C3 gene expression by interferon and STAT1 pathway have been described, potentially through indirect regulation from cytokine response and interaction with NF-kB pathway.^34^

### Ahr activity in SMC protects against dioxin-induced oxidative stress, and promotes a stable plaque phenotype

To better understand the role of AHR in mediating the effect of TCDD in lesion development, we performed scRNA-seq of atherosclerotic aorta from SMC-LnT and SMC-AhrKO mice treated with HFD and TCDD for 16 weeks (**Figure 4A**). There was a significant increase in the proportion of CMC with corresponding reduction in the FMC population among the tdTomato+ SMC-lineage cells. There was no significant difference in the overall SMC population (**Figure 4B**).

**Figure 4.**
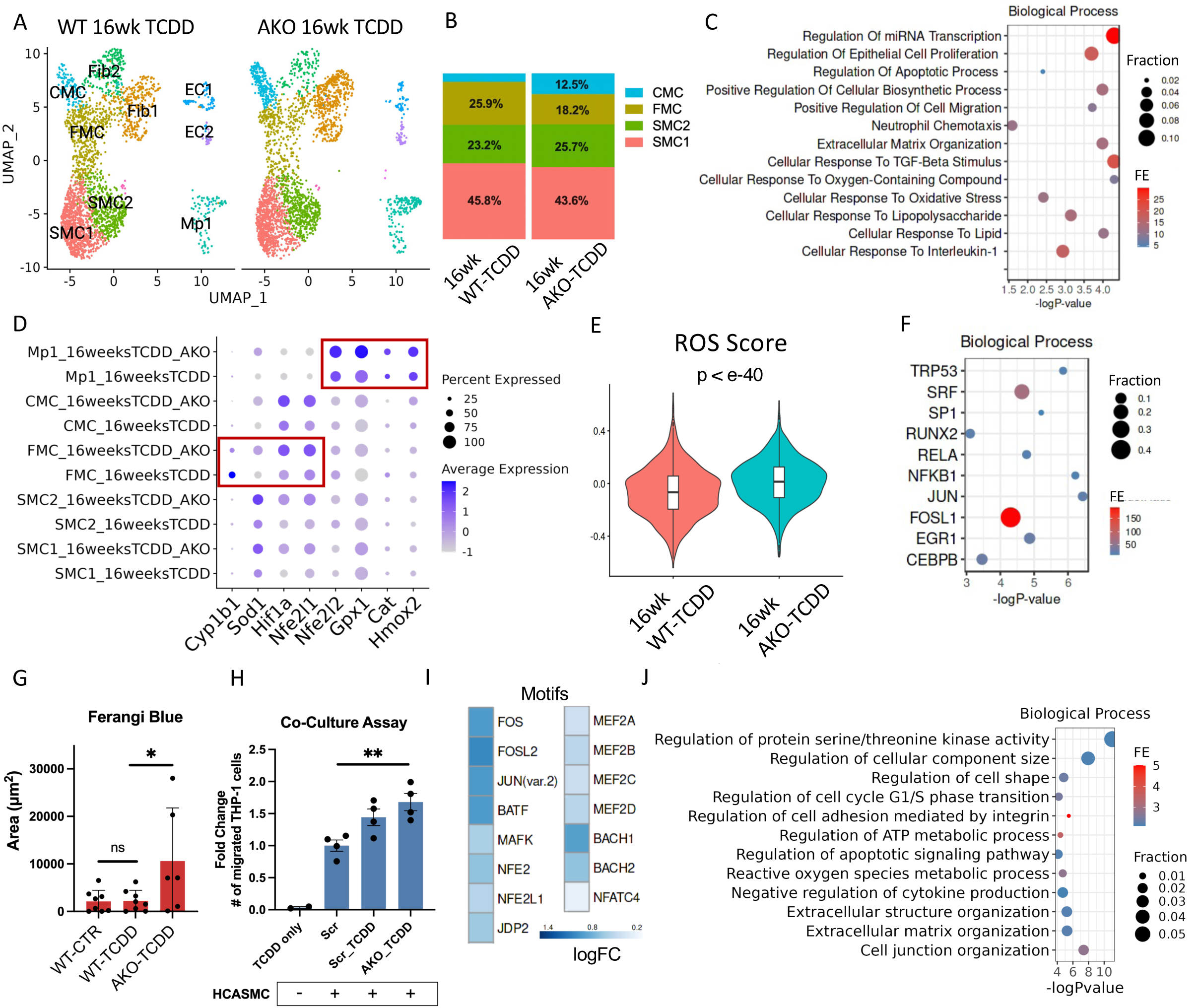
*Ahr* in SMC confers protection against dioxin-induced oxidative stress and promotes a stable plaque phenotype. (A) UMAP projection of scRNA-seq PCA clustering from SMC-LnT mice treated with 16 weeks of HFD+TCDD (n=3) or SMC-AhrKO mice treated with 16 weeks of HFD and TCDD (n=3) from 8 weeks of age. (B) A significant shift in the proportion of cells within the tdTomato+ SMC lineage cells was observed (Chi-square p < 2.2e-16). There was decrease in FMC (25.9% to 18.2%) and increase in CMC (5.1% to 12.5%) clusters. Overall, there was no significant change in the overall SMC proportions (69% to 69.3%). (C) The differentially regulated genes in FMC cluster in the SMC-AhrKO+TCDD group enrich for cell proliferation/migration/adhesion, ECM organization, cellular response to oxidative stress/TGF-beta stimulus and IL-1. (D) Oxidative stress related genes including *Sod1, Gpx1, Cat, Hmox2, Nfe2l1/Nfe2l2,* and *Hif1a* expression are increased in the SMC-lineage cells and macrophage cluster of AhrKO-TCDD group, while the *Ahr* downstream gene *Cyp1b1*, a key regulator of redox homeostasis is decreased in FMC as expected. (E) A ROS gene expression score based on average expression of the oxidative stress related genes (ros_score, *see Methods*) is overall increased in the tdTomato+ SMC-lineage cells of 16wksTCDD_AKO group compared to the 16wksTCDD group (p=2.3e-41). All subgroups of SMC-lineage cells had increased average expression of ROS genes (**Figure S4C**). (F) Upstream analysis of the upregulated genes show enrichment for transcription factors including JUN, NFKB1, TP53, SP1, and RELA, suggesting significant epigenetic activation of the pro-inflammatory and (apoptotic) pathways of AP-1, and NF-kB pathways. (G) Ferangi blue staining was performed to measure ossifying cells in the lesion. There was no change in the WT-TCDD compared to WT-CTR (p=NS), but a significant increase in ossifying region was seen in the AKO-TCDD vs. WT-CTR and WT-TCDD (p<0.05). (H) Co-culture assay with THP-1 monocytes and HCASMC show increased monocyte migration across the barrier in the setting of SMC exposure to TCDD. AHR KO resulted in further migration of THP-1 cells (ANOVA p =0.0002). Minimal migration was seen in the negative control (TCDD only, no HCASMC). (I) Differential comparison found motifs including AP-1, MAFK, NFE2/NFE2L1, MEF2, BACH1/2 to be enriched in the open chromatin regions of AKO-TCDD. (J) Enrichment analysis for gene activity based on proximity to differential chromatin accessibility found regulation of cell shape, cell adhesion, extracellular matrix organization, ROS metabolic process, and regulation of cytokine production.

The differentially regulated genes in the FMC cluster of AKO-TCDD compared to WT-TCDD enriched for pathways including cell proliferation and migration, ECM organization, response to oxidative stress, IL-1 and TGF-b stimulus (**Figure 4C, Table S6**). The average C3 expression was lower in the AKO-TCDD FMC population compared to WT-TCDD suggesting that Ahr-dependent dioxin response may partially activate the C3 pathway (**Figure S4B**). Genes related to oxidative stress were upregulated including *Gpx1, Hif1a, Sod1, Nfe2l1* in the SMC lineage cell clusters as well as macrophages (**Figure 4D, 4E**). Upstream analysis of the upregulated genes in the AKO-TCDD group of FMC found enrichment for transcription factors including JUN, NFKB1, RELA, TP53, SRF and SP1 (**Figure 4F**). These single-cell findings suggest that SMC-specific loss of *Ahr* leads to increased oxidative stress and inflammation resulting in increased lesion size and macrophage content.

Furthermore, consistent with the increase of CMC in the single-cell data, we found the relative alkaline phosphatase (AP)-stained area to be increased in the AKO-TCDD cohort compared with WT-CTR and WT-TCDD groups (P<0.05, **Figure 4G**).

There was also increase in Tagln+ area in lesion likely reflecting the overall increase in lesion size (**Figure 3D**). The histological assessment for C3 protein expression showed decreased signal in the lesion of AKO-TCDD, which likely reflect the reduction in the proportion of C3 expressing FMC in the AKO-TCDD group (**Figure 3K**). Overall, these findings in the AKO-TCDD group, consisting of increased lesion size and inflammatory burden, and reduced SMC content in lesion cap are reflective of a more vulnerable plaque phenotype.

### Ahr knockout in SMC increases monocyte migration and lead to significant epigenetic activation of inflammatory pathways in SMC-lineage cells

We further assessed the cellular cross-talk between dioxin-induced inflammatory SMC and macrophages in a co-culture model utilizing human coronary artery smooth muscle cell (HCASMC) and THP-1 cell lines *in vitro*. We found a significant increase in migration of monocyte/macrophages following HCASMC exposure to dioxin. Minimal migration was seen in the absence of HCASMC. AHR KO in HCASMC resulted in further increase in migration of monocytes (**Figure 4H**). These findings are consistent with the scRNA-seq data showing increased proportion of macrophages in the AKO-TCDD compared to WT-TCDD, and the increased Cd68+ area in the histological analysis (**Figure S4C**). We then explored if there were differences in M1/M2 polarization of macrophages using well-known markers of M1(pro-inflammatory)- and M2(anti-inflammatory)-like macrophages.^35, 36^ We found that the M1 markers were upregulated, and M2 markers were downregulated in the AKO-TCDD group macrophages compared to WT-TCDD consistent with pro-inflammatory changes (**Figure S4D**).

To understand how loss of AhR in SMC affects the epigenetic response to TCDD, we mapped the chromatin accessibility landscape using scATAC-seq in AKO-TCDD group compared to the WT-TCDD group. We found a significant number of differentially regulated peaks in the FMC and SMC clusters (**Table S7**). The upregulated accessible (DA) chromatin peaks in FMC enriched for motifs for TFs related to inflammatory and oxidative stress including AP-1, BACH1, MAF/NFE2L1/NFE2 and NF-κB/RELA (**Figure 4I**). The DA peaks in FMC enriched for genes related to biological pathways including regulation of cell shape, cell adhesion, extracellular matrix organization, ROS metabolic process, and regulation of cytokine production (**Figure 4J**). The DA peaks in SMC enriched for motifs JUN/FOS, BACH1, NFE2L1, and REL/RELA, and these peaks enriched for genes related to cell proliferation, migration and blood vessel development (**Figure S4E, S4F**).

## Discussion

Dioxin is a major environmental pollutant prevalent in the ecological system and is a component of several major cardiovascular risk factors, including tobacco smoke and air pollution. Epidemiologic data indicate that exposure to dioxin and dioxin containing compounds is associated with increased cardiovascular morbidity and mortality.^18–20, 37^ These studies collectively highlight the significant public health impact of dioxin exposure on cardiovascular disease development. Dioxin induces oxidative stress and inflammation in various tissue types and disease environments, largely mediated through activation of the AHR pathway, a ligand-activated bHLH/PAS transcription factor.^38, 39^ The promiscuity of AHR results in its activation by various environmental toxicants, including dioxin, polycyclic aromatic hydrocarbons, and acrolein, which are commonly present in cigarette smoke and air pollution. In the context of atherosclerosis, dioxin exposure leads to increased disease burden in mouse models, associated with heightened inflammatory biomarkers, circulating inflammatory cells, and the accumulation of oxidized lipids within atherosclerotic lesions.^30, 40^

In this study, we identified significant transcriptional and epigenetic effects of dioxin on vascular smooth muscle cells (SMC), leading to their phenotypic modulation and increased atherosclerotic burden. Dioxin appears to promote disease through two main mechanisms: (i) modulation of SMC phenotype in response to vascular injury and (ii) induction of a pro-inflammatory state, upregulating the complement pathway in SMC-lineage cells, which contributes to increased macrophage recruitment. Moreover, in SMC-specific Ahr knockout (KO) models, we observed increased markers of oxidative stress and features indicative of plaque instability, suggesting that *AHR* is protective in modulating dioxin-induced vascular disease.

Our findings are novel, demonstrating a mechanistic link between environmental toxicants and vascular disease, with *AHR* playing a critical role in SMCs. Single-cell analyses and in vitro validation studies revealed that dioxin induces early SMC de-differentiation, characterized by increased expression of FMC markers, and promotes SMC proliferation and migration. Concurrently, we observed heightened expression of inflammatory genes in the SMC lineage cells, including NF-kB and innate immunity pathways, such as complement activation.

Further analysis revealed that dioxin activates the complement pathway within SMCs, which likely drives macrophage recruitment. We observed upregulation of complement receptors (*Itgam/Cd11b, Itgb2, C3ra1*) and Toll-like receptor 4, both of which are key regulators of innate immune response. Complement activation has been shown to enhance macrophage recruitment in atherosclerosis and other inflammatory diseases, and has been proposed as a signature of an inflammatory SMC phenotype leading to high level of cytokine production.^41, 42^ Furthermore, oxidized lipids and lipoproteins have been reported to act as stimulatory trigger for complement in lesions.^43^ C3, a central component of the complement cascade, was also found highly expressed in clonally expanding SMC in atherosclerosis,^44^ suggesting that dioxin may promote the clonal expansion of *C3*+/*Sca1*+ modulated SMC, which may further drive the inflammatory milieu within atherosclerotic lesions.^45^ The acceleration of clonal expansion of SMC mediated by dioxin represent a potentially interesting connection with dioxin’s well-known tumorigenic properties and warrant further investigation. These findings point to the *C3*+ modulated-SMC as a possible target for therapy and a pathway by which dioxin promote atherosclerosis. The scATAC-seq analysis of modulated SMC also found transcription factors of inflammation activated following TCDD exposure including the IRF family and JAK-STAT pathways, which are critical factors in regulating innate immunity and complement activation.^46^

Consistent with these findings, the histological analysis of the atherosclerotic lesion from mice exposed to long-term dioxin resulted in increased lesion size, increased area of tdTomato+ SMC lineage cells and macrophages in the lesion. The increase in the macrophage content and lesion size are consistent with previously reported findings by Wu et al.^30^ Contrary to the short-term exposure data, snap shots of the aortic atherosclerotic lesion after 8 and 16 weeks of TCDD treatment using the SMC-lineage mouse model found a decrease in the proportion of modulated SMC. Correlating these findings with the histological observations of increased tdTomato+ SMC lineage cells in the lesion and lesion cap, the single-cell data is reflective of modulated SMCs differentiating back to the quiescent SMC state in the lesion following prolonged exposure to TCDD. These findings suggest that dioxin likely accelerates the natural atherosclerotic process including SMC phenotypic modulation to create the protective lesion cap. The striking reduction in global TCF21 occupancy in HCASMC suggest a significant epigenetic mechanism underlying the SMC phenotypic response to dioxin exposure. Further studies of key SMC transcription factors and their response to dioxin is warranted.

*AHR* plays a complex role in oxidative stress regulation, balancing both pro-oxidative and anti-oxidative responses depending on the specific cellular context.^47–49^ Moreover, *AHR*-independent effects of dioxin and dioxin-independent functions of *AHR* have been reported.^27, 50^ In this study, we further assessed the role of *AHR* of SMC in modulating the dioxin effect on atherosclerosis. We assumed that if the dioxin effect on SMC was predominantly driven by *AHR* pathway activation, then the dioxin exposure SMC phenotype would reverse in the *Ahr* KO model. To the contrary, we observed further increase in the lesion content of SMC-lineage cells and macrophages in AKO-TCDD compared to WT-TCDD. In addition, the tdTomato+ cells were significantly absent from the fibrous cap area in the AKO-TCDD model, suggesting that SMC failed to reach the cap or they traveled back into the plaque. These results are consistent with our previous finding demonstrating the critical role of *AHR* in maintaining functional SMC phenotypic modulation.^27^

The increase in lesion size in the AKO-TCDD group is consistent with the increased oxidative stress and pro-inflammatory signal seen in the scRNA-seq and scATAC-seq compared to the WT-TCDD group. These analyses suggest that the lack of *Ahr* in SMC results in a pro-inflammatory state of SMC-lineage cells, leading to release of paracrine factors increasing recruitment of macrophages. Interestingly, despite the overall increase in inflammatory phenotype, the *C3* expression level in the modulated SMC was lower in the AKO-TCDD compared to WT-TCDD (**Figure S4B**). Despite the mild decrease in C3 expression in AKO-TCDD, there was overall increased lesion size and macrophage infiltration into the lesion, suggesting that *Ahr* KO results in overall pro-inflammatory milieu. These findings were supported by results from migration assays with monocytes and HCASMC showing further increase in dioxin-induced macrophage recruitment when *AHR* is knocked out from SMC. Our results demonstrate that *AHR* in SMC exhibit both pro- and anti-inflammatory activity in response to dioxin with a net protective role in dioxin-induced atherosclerosis.

The current data provides further evidence that *AHR* in SMC is a key regulatory gene of a protective vascular remodeling in response to atherosclerotic pro-inflammatory insult, both by providing an anti-oxidative response and promoting lesion stabilization through the formation of SMC-rich lesion cap. One interesting observation is that when comparing the phenotype of the previously published *Ahr* KO mice phenotype to the current study, the Cd68+ macrophage proportions are similar between the *Ahr* KO group and WT in the absence of dioxin.^27^ This suggest that the presence of dioxin, a pro-oxidative stress inducer, appears to bring out the gene-environment interaction resulting in further increase in the macrophage recruitment to the lesion.

This study has several limitations – we characterize the effect of dioxin with focus on the SMC lineage cells. Exposure to dioxin is not limited to the SMCs and we find that there is significant dioxin response in other cell types including fibroblasts and endothelial cells. The impact of dioxin on the other cell types including macrophages and endothelial cells have been described before, and those of the vascular wall should be studied further with appropriate models to validate the findings. Some of the findings from this study are exploratory – based on cell-cell interaction analysis based on scRNA-seq, we propose a mechanism by which SMC becomes pro-inflammatory and secretes cytokines and complement factors to attract macrophage into the lesion. These findings support previous observations that dioxin increase the inflammatory burden of atherosclerotic lesion, through both Ahr-dependent and independent pathways. The protective function of Ahr is consistent with previous reports of pro- and anti-inflammatory function of AHR which is context and cell-type specific.

## Conclusion

In conclusion, this study demonstrates that dioxin exposure accelerates atherosclerosis by promoting SMC phenotypic modulation, inflammation, and oxidative stress through AHR-regulated pathways. SMCs exposed to dioxin take on a pro-inflammatory role, increasing the secretion of complement factors and cytokines, thereby enhancing macrophage recruitment and contributing to plaque development. Notably, the absence of AHR in SMCs exacerbates the atherosclerotic phenotype, resulting in increased oxidative stress, inflammatory signaling, and a more vulnerable plaque composition. These findings underscore the protective role of AHR in stabilizing atherosclerotic lesions and regulating SMC behavior under dioxin exposure (**Figure 5**). By elucidating the molecular mechanisms linking dioxin exposure to vascular disease, this study highlights potential therapeutic targets, including AHR and the C3+ modulated SMC population, which could pave the way for strategies to mitigate environmental contributions to cardiovascular disease.

**Figure 5.**
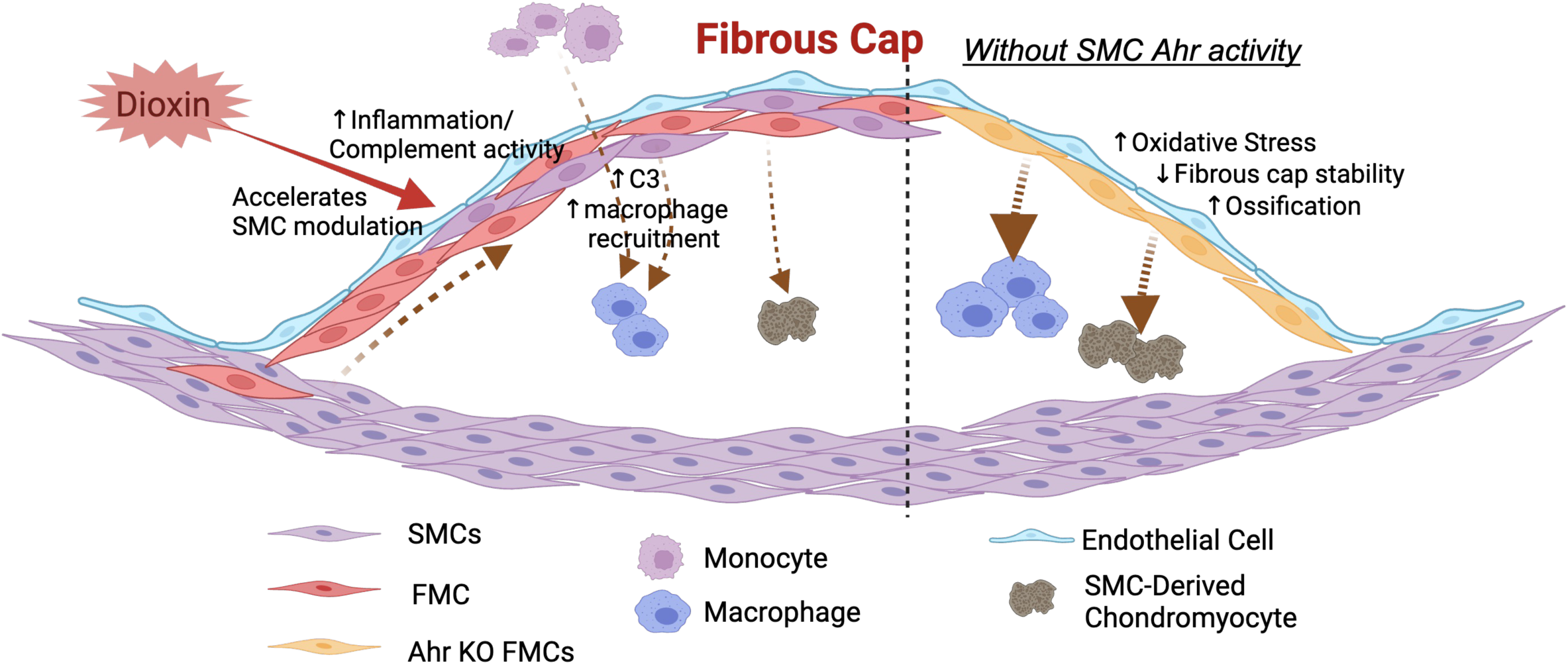
Dioxin accelerates atherosclerosis and AHR in SMC is protective in mediating response to dioxin (and AHR loss further accelerate SMC response to dioxin) Dioxin i) accelerates the SMC modulation during lesion development resulting in increased SMC migration to lesion cap and lesion intima; ii) increase the inflammatory response of SMC resulting in cytokine production and activation of the complement pathway, leading to increase in macrophage recruitment to the lesion. Loss of Ahr in SMC-lineage cell results in a more vulnerable plaque phenotype including loss of SMC from the protective lesion cap, increased oxidative stress, macrophage recruitment and lesion size.

