## Supplemental Materials for "Aryl-hydrocarbon receptor in smooth muscle cells protect against dioxin induced adverse remodeling of atherosclerosis"

**Supplemental Table S1-S7.**

Table S1. Differentially expressed genes following short-term TCDD exposure in DRE-LacZ Tg mice scRNA-seq (FMC, SMC1/2, MP1, Fib2)

Table S2. Differentially expressed genes with dioxin treatment (bulkRNAseq).

Table S3. Differentially expressed genes with CSE treatment (bulkRNAseq).

Table S4. Differentially expressed genes with 8 weeks TCDD treatment in SMC, FMC and CMC (scRNAseq).

Table S5. Differential accessibility (DA) peaks following TCDD exposure in FMC (scATACseq).

Table S6. Differentially expressed genes between WT-TCDD and AKO-TCDD (16 weeks TCDD treatment in both groups)

Table S7. Differential accessibility (DA) peaks in AKO-TCDD group compared to the WT-TCDD in FMC and SMC (scATACseq).

#### Supplemental Figure Legends

**Figure 1.** A. Histology of atherosclerotic aortic sinus from DRE-lacZ Tg mouse The lacZ signal is present in the medial layer and lesion cap ; ApoE KO mouse exposed to dioxin show β-gal staining in the media and lesion cap (blue arrow). B. Featureplot depicting gene expression of SMC marker gene (Cnn1), and modulated SMC marker genes (*Lum, Vcam1, Spp1*). C. There is significant increase in Cyp1b1 expression in the modulated SMC population following TCDD exposure. D. Inflammatory gene expression is increased following TCDD exposure in Mod-SMC and Fibroblast population. E. There is increase in oxidative stress as measured by reactive oxygen species following TCDD exposure ($p=0.053$). F. Cell-cell interaction analysis with *CellChat* found SPP1 signaling pathway with significant outgoing influence from Mod-SMC following TCDD exposure, and identifies Mod-SMC as key mediator and influencer cell type responsive to TCDD. G. Featureplot of C3 and Itgam expression level in ctrl and tcdd groups. C3 is expressed primarily in the Mod-SMC and Fibroblast population and Itgam expression is limited to the Macrophage population.

**Figure 2.** A. RNAseq of HCASMC treated with (*left*) control (WT_rep1/2/3/4) and TCDD (D_rep1/2/3/4); (*right*) control (D2/3) and CSE (CSE1/2/3) show significant change in transcriptional profile. B. Upstream analysis of the differentially regulated genes in response to TCDD are predicted to be regulated by key SMC TFs and inflammatory TFs. C. GO Biological pathways identified as enriched based on HCASMC response to CSE include cellular response to chemical/cytokine stimulus, ER stress/protein targeting to ER, circulatory system development, oxidate stress induced cell death, and celluar localization. D. There is significant overlap in the DE genes (FDR $<0.05$, Fold change $>1.3$) of CSE and TCDD exposure ($p<e\text{-}15$). E. The upregulated chromatin accessible peaks in response to TCDD enriched for pathways including cellular response to stress, vasculature development, response to cytokine stimulus, and ECM organization. F. There was a significant overlap between the upregulated chromatin accessible regions between TCDD and CSE treatment ($p<e\text{-}255$). G. The upregulated chromatin accessible peaks in response to CSE enriched for pathways including vasculature development, regulation of cell migration, angiogenesis, MAPK/NF-kB signaling, ER stress and apoptosis. H. There is overall increase in chromatin accessibility following CSE exposure (blue) and TCDD (red) exposure around shared genomic locations. I. (left) Using *macs2 bdgdiff* analysis, there is an overall increase in AHR binding on ChIP-seq following TCDD treatment (left, blue denotes increased binding), (right) There is clear decrease in binding of TCF21 on ChIP-seq, centered around the AHR binding peaks in the presence TCDD.

**Figure 3.** A. Schematic of the animal exposure and tamoxifen/HFD treatment experiment. B. Featureplot of the SMC marker (Cnn1), FMC marker (Lum), CMC marker (Spp1), and tdTomato expression in the WT (16 wks HFD) and 8 weeks TCDD (16wks HFD) combined umap. C. Cyp1b1 expression is localized to the FMC and Fibroblast populations, and increases with TCDD treatment. D. Significant transcriptional changes occur in FMC population following TCDD as visualized by the volcano plot. E. Oxidative stress related genes are increased following TCDD exposure in SMC-lineage cell types. F. Representative immunohistochemistry images of Tagln are shown. G. Immunostaining for Itgam show increase following TCDD treatment, however, no significant change in the AKO-TCDD group. (ANOVA p=NS). H. Markers of M1-like polarization of macrophage were used to construct a “M1 polarization” gene expression score based on average expression of genes (*AddModuleScore* in Seurat, *See Supplemental Methods*). There is significant increase in the M1 polarization signal in the macrophage population ($p<e-77$). I. Immunostaining for C3 and Itgam show adjacent localization of the tdT+/C3+ cells and the Itgam+ cells (10x magnification). J. (left) Motifs including MEF2A/B/C/D, TEAD1/2/3/4, CTCF, and TCF3/12 are significantly enriched in the SMC population after TCDD treatment (right) Motifs including ETV5/6, ELK1/2/3/4, IRF1/7/8/9, CTCF, and TCF12 are significantly enriched in the macrophage population after TCDD treatment.

**Figure 4**. A. UMAP of scATAC-seq results show the different treatment groups WT (WT-CTR), TCDD (WT-TCDD) and TCDDAKO (AKO-TCDD). The different SMC-lineage cell states are identified based on label transfer by anchoring to scRNA-seq data. B. No significant difference was found in the C3 expression levels between WT-TCDD and AKO-TCDD groups in FMC and CMC. Expression level was very low or absent in SMC. C. There is shift in tdTomato-negative cell proportions in AKO-TCDD compared to WT-TCDD, including increased macrophage and Fib1 population (Chi-square $p<2.2e-16$). D. There is shift in tdTomato-negative cell proportions in AKO-TCDD compared to WT-TCDD, including increased macrophage and Fib1 population. E. Motifs including JUN/FOS (AP-1), MAFK, BACH1/2, BATF, NFE2L1, RELA are significantly enriched in the SMC population of AKO-TCDD to WT-TCDD. F. Differentially accessible peaks in AKO-TCDD compared to WT-TCDD enrich for biological pathways including cell migration, cell proliferation/differentiation, and blood vessel development.

### Supplemental Figure 1.

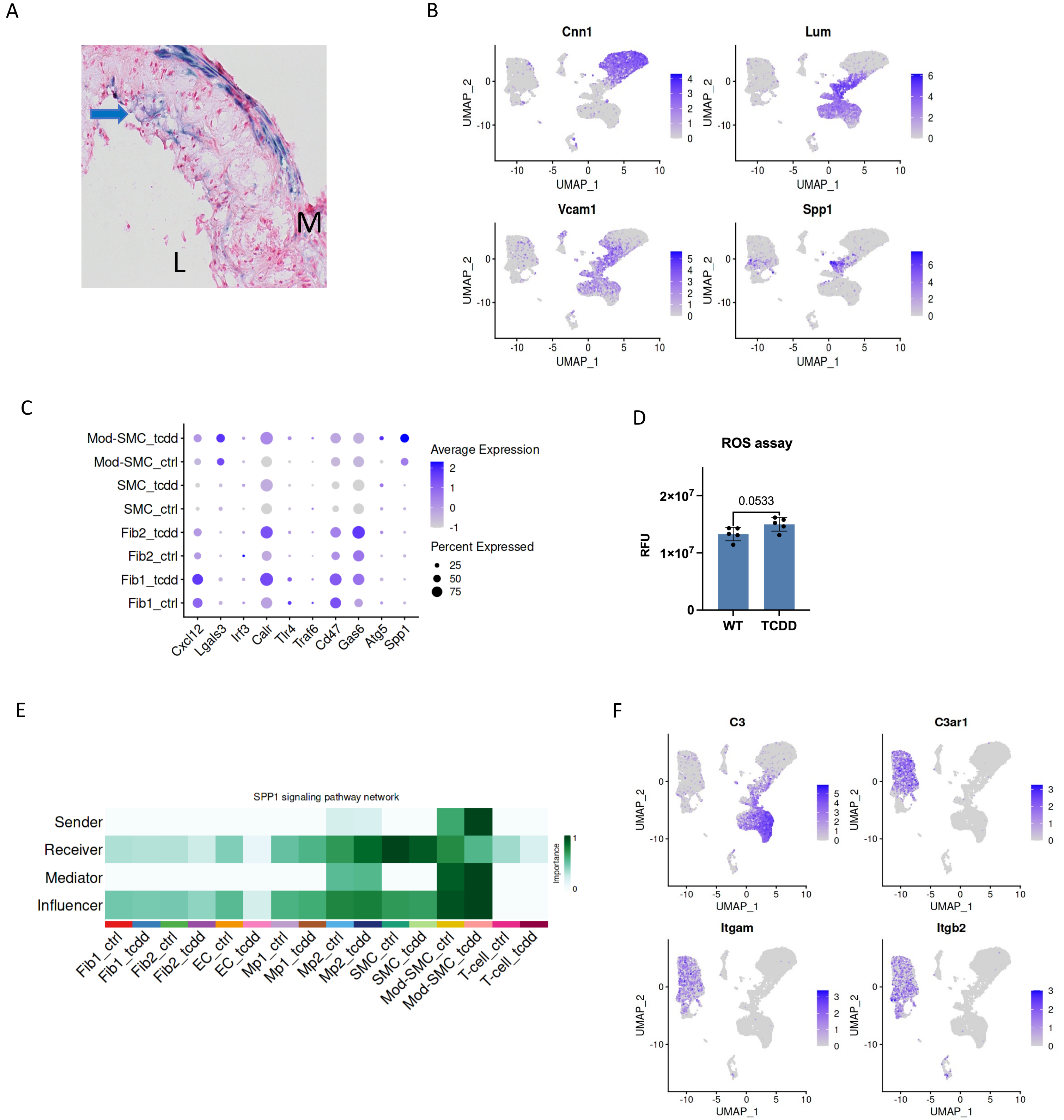

### Supplemental Figure 2.

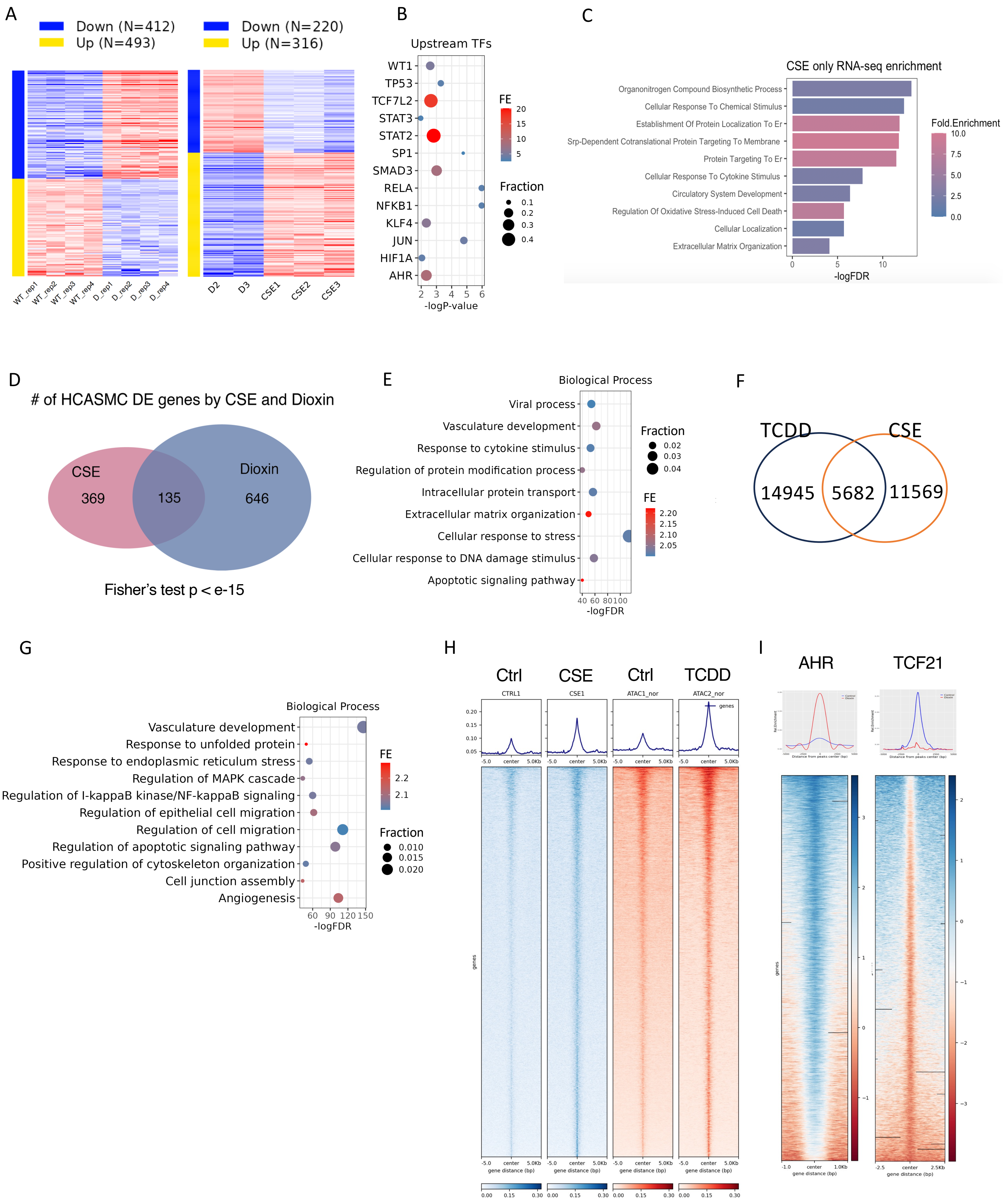

### Supplemental Figure 3.

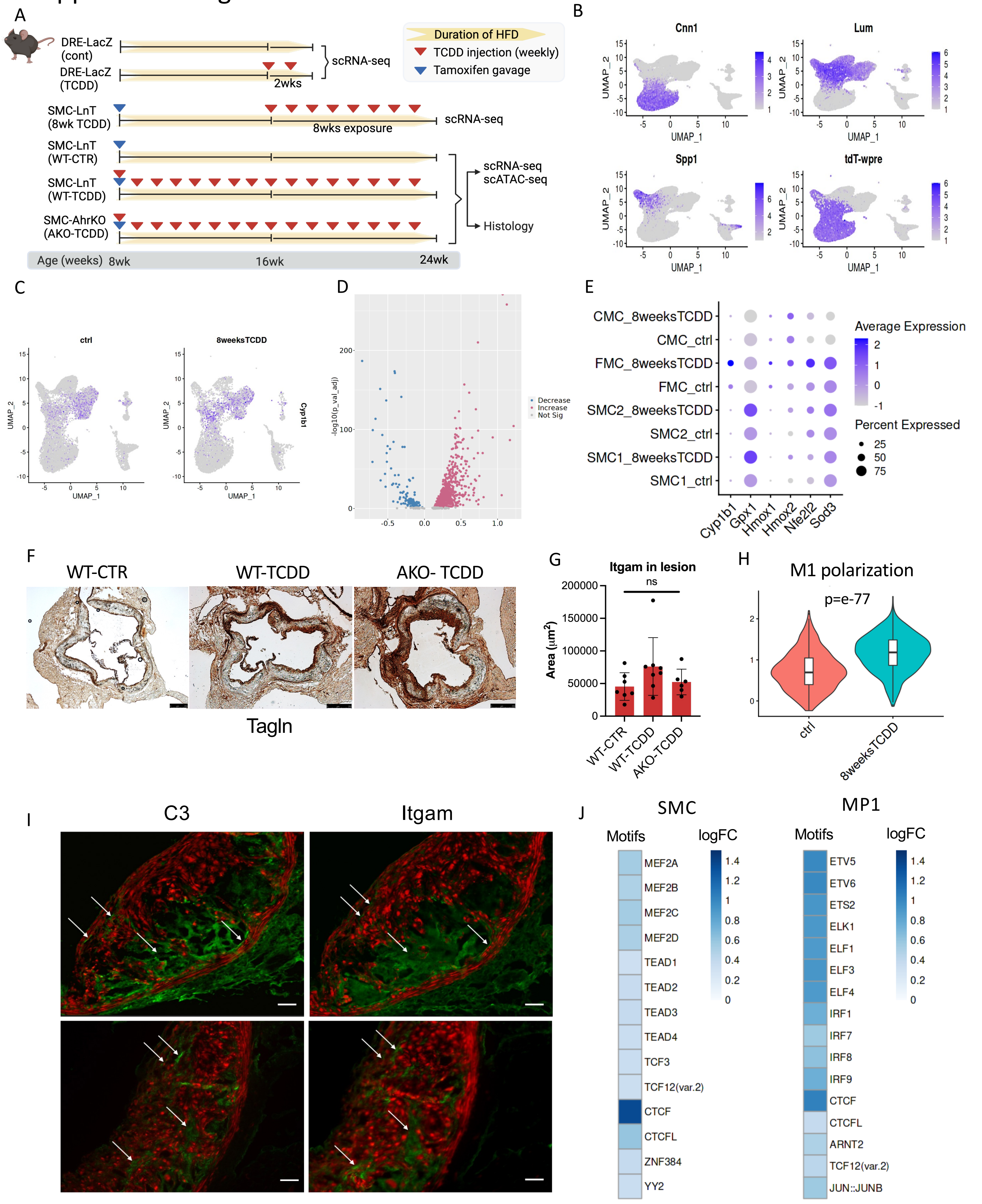

### Supplemental Figure 4.

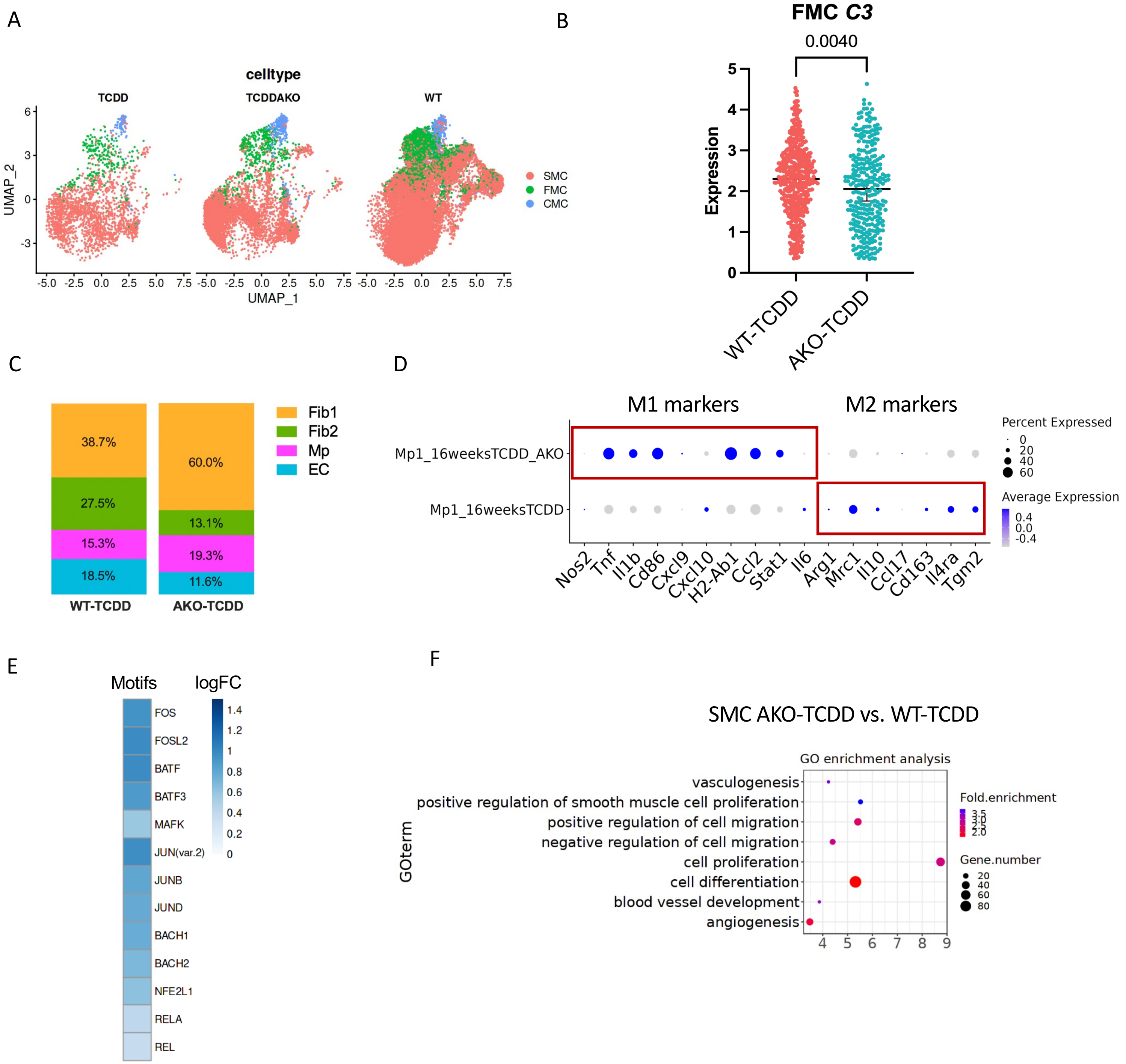

#### Supplemental Methods

##### Chemicals

The 2,3,7,8-Tetrachlorodibenzo-p-dioxin (TCDD) was purchased from AccuStandard Inc (D-404N) and dissolved in DMSO (Thermo Scientific, S7563) to create a 770 µM stock solution. Cigarette smoke extract (CSE) was generated using the Scireq inExpose system (Montreal, QC, Canada) with the Flexiware software (v8.0). The system was configured with a cigarette smoking robot (SCIREQ) for automated delivery of cigarette smoke to a flask containing 20 mL of complete cell culture medium with 5% serum. Specifically, one cigarette, approximately nine puffs, was smoked into cell culture medium according to the following parameters (ISO 1991): one 35 mL puff of 2-second duration with a puff interval of 60 seconds. MTT assay was performed to assess the non-cytotoxic range of concentration.

##### Culture of Primary HCASMC, immortalized HCASMC and THP1 Cell

Primary human coronary artery smooth muscle cells (HCASMC) were utilized for in vitro experiments in this study. Cells were purchased from Cell Applications, Inc (San Diego, CA). We used smooth muscle growth medium (SmBm) (Lonza, CC-3182) for cell culture. All HCASMC lines were used between passage 5-8. The immortalized hTERT HCASMC were obtained from Clint Miller lab. In general, the HCASMC were purchased from Cell Applications (Catalog No. 350-05a) and immortalized by lentiviral transduction of the hTERT-IRES-hygro construct (Addgene Plasmid, 85140) in 10 µg/ml polybrene. Transduced cells were selected using 400 µg/ml of hygromycin (Gibco, 10687010) and maintained in SmBm medium for cell culture. The THP-1 cell line was purchased from company. THP-1 cells were cultured in RPMI 1640 medium (Corning, 15040CV) supplemented with 10% FBS (Sigma-Aldrich, 12306C).

##### RNA isolation and qRT-PCR

RNA was isolated using RNeasy mini kit (Qiagen, 74104) and reversely transcribed to cDNA using High-Capacity RNA-to-cDNA kit (Thermo Fisher, 4387406). Gene expression levels were measured using Taqman probes (Invitrogen) (LUM, IL1A, CNN1, ACTA2, CYP1B1) for SYBR Green assay and quantified on a ViiA7 Real-Time PCR system (Applied Biosystems, Foster City, CA) and normalized to GAPDH levels.

##### HCASMC phenotypic assays

Migration and proliferation of HCASMC was quantified as described previously.[1] The Radius Cell Migration Assay kit (Cell Biolabs) was used for the gap closure assay. HCASMC was plated at 80% confluence, then following the protocol, the polymer at the center of each well were dissolved to allow migration. Migration was quantified by remaining area after migration, compared to origin area. For the proliferation assay, 5-ethynyl-2′-deoxyuridine was introduced into the cell culture 3 hours before assay for uptake. The protocol for Click-iT Plus 5-ethynyl-2′-deoxyuridine proliferation kit (Thermo Fisher) was followed as instructed.

##### Oxidative stress assays

To test the oxidative stress, Reactive Oxygen Species (ROS) formation in HCASMC cells was detected using a DCFDA/H2DCFDA-Cellular ROS Assay Kit (Abcam, ab113851) according to the manufacturer’s protocol. Generally, 25,000 cells were seeded per well in 96-well microplate and cells were allowed to adhere overnight. Then cells were stained with diluted DCFDA solution for 45 minutes at 37℃ in the dark followed by treated cells with TCDD or vehicle for 24 hours. Finally, the microplate was measured on a fluorescence plate reader (SpectraMax iD3) at Ex/Em = 485/535nm in end point mode.

##### Macrophage migration assay

The hTert HCASMC were grown to 70% confluence in 24-well plate, then transfected with AHR siRNA or scramble control to final concentration of 20nM with RNAiMax (Invitrogen, 13778075) for 24 hours. The small interfering RNA (siRNA) for AHR was purchased from Origene (SR300136). Then change to SmBm medium for 24 hours, followed by 10nM TCDD treatment or SmBm control for 24 hours. Then a cell migration assay was performed to study the migration behavior of THP-1 macrophage cells. Cell culture inserts for 24-well plate containing a membrane with pores of 8 µm (Corning, 353097) were used. Fifteen thousand THP-1 macrophages in 200 µL of 1% FBS RPMI-1640 were loaded into both inserts and 24-well plates. Control cells were treated with the vehicle alone (DMSO). The transwell chambers were then incubated at 37 °C in 5% (v/v) $CO_2$ for 24 h to evaluate cell migration. Then, the inserts were removed and THP-1 cells that migrated through the membrane to the lower chamber was calculated based on images.

**RNA sequencing and analysis**
Primary HCASMC were grown to 70% confluence in 6-well plate. The second day, cells were treated with 10nM TCDD, CSE or vehicle control for 24 hours. Four replicates were used for each condition for TCDD RNAseq. Three replicates were used for each condition for CSE RNAseq. RNAs were sent to Novogene for sample QC, library preparation, and sequencing. All samples passed QC, and 250-300 bp insert cDNA libraries were prepared for each sample. Subsequently, sequencing was performed on a Novaseq 6000 platform with paired-end 150 bp reads. The script for RNA-Seq analysis can be found in GitHub (https://github.com/zhaoshuoxp/Pipelines-Wrappers/blob/master/RNAseq.sh). Additionally, the web-based tools iDEP 2.01 (http://bioinformatics.sdstate.edu/idep/) was used for analysis of the RNA-Seq data and visualization using the counts data generated from FeatureCounts.[2]

**ChIP-Seq assay and analysis**
ChIP was performed using the AHR antibody (Santa Cruz, sc-5579) and TCF21 antibody (HPA013189, Sigma), which were pre-validated according to ChIP-seq guidelines. Two replicates were used for each condition. Library for ChIP-Seq was prepared using standard procedures as described previously.[3] Briefly, approximately 4,000,000 HCASMC treated or not treated with TCDD were fixed with 1% formaldehyde and quenched by glycine. The cells were washed three times with PBS and then harvested in ChIP lysis buffer (50 mM Tris-HCl, pH 8, 5 mM EDTA, 0.5% SDS). Cross-linked chromatin was sheared for 3 × 1 min by sonication (Branson SFX250 Sonifier) before extensive centrifugation. Four volume of ChIP dilution buffer (20 mM Tris-HCl, pH 8.0, 150 mM NaCl, 2 mM EDTA, 1% Triton X-100) was added to the supernatant. The resulting lysate was then incubated with DynabeadsTM Protein G (Thermo Scientific, 10009D) and antibodies at 4 °C overnight. After bead washing, DNA was eluted by ChIP elution buffer (0.1 M NaHCO3, 1% SDS, 20 µg/ml proteinase K). The elution was incubated at 65 °C overnight, and DNA was extracted with a DNA purification kit (Zymo, D4013). The purified DNA was assayed by quantitative PCR with ABI ViiA 7 and Power SYBR Green Master Mix (ABI, 4368706) using CYP1B1 DRE region primers. Library for ChIP-Seq was prepared using standard procedures. Briefly, Libraries were prepared with KAPA Hyper Prep kit (KK8502). ChIPseq libraries were sequenced on HiSeq X10 for 150 bp paired-end sequencing. The script for RNA- Seq analysis can be found in GitHub (https://github.com/zhaoshuoxp/Pipelines-Wrappers/blob/master/ChIPseq.sh). Briefly, quality control of ChIPseq data was performed using *Fastqc*, and then low-quality bases and adaptor contamination were trimmed by *cutadapt*. Filtered reads were mapped to hg19 using BWA mem algorithm. Duplicate reads were marked by *Picard Markduplicate* module and removed with unmapped reads by *samtools* view *-f 2 -F 1804*.

*macs2 callpeak* was used for peaks calling and input as control. *macs2 bdgdiff* was used for differential peaks calling with default parameters. The regions specific to each sample (cond1.bed and cond2.bed) were intersected with the peaks identified in each sample using *macs2 callpeak*. Peaks in each sample that overlapped with the regions identified by *bdgdiff* were classified as differential peaks. Peaks that overlapped between the two samples and did not overlap with differentially identified regions from bdgdiff were categorized as common peaks. Bigwig files were generated for Genome Browser visualization.

For enrichment level analysis, reads of control/treatment samples which mapped on peaks were counted by *intersectBed* and normalized by per million (RPM). ChIPseq peaks, or open chromatin regions were clustered using the hierarchical algorithm, and reads centering on these peaks (± 5 kb) were plotted with *deeptools*.

We utilized the *Genomic Regions Enrichment of Annotations Tool (GREAT 4.0)* to analyze the detected peaks, with the parameter "single nearest gene," which is within 50 kb to nearest genes.[4] Gene ontology from *GREAT* output was analyzed by *DAVID*. *KEGG* pathways and biological processes enrichment analysis was carried out using default settings. The *HOMER findMotifsGenome*.pl script was employed to search for known *TRANSFAC* motifs and to generate *de novo* motifs.[5]

**Assay of transposase-accessible chromatin (ATAC) and analysis**
For TCDD ATAC-Seq, we followed the previous protocol.[6] Two replicates were used for each condition. Approximately 5e4 fresh HCAMSC cells were collected by centrifugation at 500 g and washed twice with cold PBS. Nuclei-enriched fractions were extracted with cold resuspension buffer (0.1% NP-40, 0.1% Tween 20, and 0.01% Digitonin) and washed out with 1 ml of cold resuspension buffer containing 0.1% Tween 20 only. Nuclei pellets were collected by centrifugation and resuspended with transposition reaction buffer containing Tn5 transposases (Illumina Nextera). Transposition reactions were incubated at 37 °C for 30 min, followed by DNA purification using the DNA Clean-up and Concentration kit (Zymo, D4013). Libraries were amplified using Nextera barcodes and high-fidelity polymerase (NEB, M0541S) and purified using Agencourt Ampure XP beads (Beckman Coulter, A63880) double-size selection (0.5X:0.9X). Libraries were sequenced on HiSeq X10 for 150bp paired-end sequencing. The CES ATAC-seq was performed following the manufacturer's instructions (Active Motif, 53150). Generally, 100,000 HCASMC were collected and lysed in the ATAC-seq lysis buffer. Next the samples were processed for the transposase reaction, library generation and purification. The samples were sent to Novogene and sequenced.

Data were analyzed as described previously.[3] The script for ATAC-Seq analysis can be found in GitHub (https://github.com/zhaoshuoxp/Pipelines-Wrappers/blob/master/ATACseq.sh). Libraries were sequenced on HiSeq X10 for 150bp paired-end sequencing. Raw fastq files were evaluated with *fastqc*, and then low-quality bases and adaptor contamination were trimmed by *cutadapt*. Reads were mapped to hg19 using bowtie2. Duplicate reads were marked by *Picard Markduplicate* module and removed with unmapped or mitochondrial reads by *samtools*. *bedtools* was used to generate BED file from filtered reads followed by Tn5 shifting with *awk*. *macs2* callpeak with *--broad* parameter was used for peak calling. *macs2 bdgdiff* with FDR cutoff 0.05 were used for differential peak comparison in AHR disrupted samples. Bigwig files were generated for Genome Browser visualization. The enrichment analysis, downstream target identification, and upstream motif analysis were conducted in a manner analogous to the analyzing of ChIP-seq data. For qPCR experiments, the purified DNA was quantified with ABI ViiA 7 and Power SYBR Green Master Mix (ABI, 4368706) and normalized by genomic DNA which extracted using Quick-DNA Microprep Kit (Zymo, D3020). Assays were repeated at least three times. Data shown were average values ± SD of representative experiments.

**Mouse strains**

The animal study protocol was approved by the Administrative Panel on Laboratory Animal Care at Stanford University. The DRE-LacZ strain was rederived from cryopreserved embryos at the Jackson Laboratories (B6.Cg-Tg(DRE-lacZ)2Gswz/J strain, JAX No. 006229) then bred to ApoE$^{-/-}$ background. SMC-specific lineage tracing and Ahr gene knockout in the atherosclerotic model was performed as described previously.[1] Generally, we used mice containing a well-characterized BAC transgene that expresses a tamoxifen-inducible Cre recombinase driven by the SMC-specific Myh11 promoter (Tg$^{Myh11-cREert2}$, JAX No. 019079). These mice were bred with a floxed tandem dimer tomato (tdT) fluorescent reporter line (B6.Cg-Gt(ROSA)26Sor$^{tm14(CAGtdTomato)Hze}$/J, JAX No. 007914) to allow SMC-specific lineage tracing to generate the SMC-LnT mice. All mice were bred onto the C56BL/6, ApoE$^{-/-}$ background. An Ahr$^{flox/flox}$ mice were obtained from Jackson Labs (JAX No. 006203), which was constructed by placing lox-P sites flanking the second exon of the Ahr gene. The Ahr$^{flox/flox}$ mice were then bred to the SMC-specific lineage tracing mice to generate the SMC-LnT mice. As the Cre-expressing BAC was integrated into the Y chromosome, all lineage tracing mice in the study were male.

Induction of lineage marker and Ahr knockout by Cre recombinase was performed as described previously.[1] Generally, tamoxifen was administered by oral gavage at 8 weeks of age to induce the lineage marker and Ahr knockout, followed by the initiation of a high-fat diet (Dyets No. 101511).

**Mouse aortic root cell dissociation**

Immediately after sacrifice, mice were perfused with phosphate buffered saline (PBS). The aortic root was excised and washed three times in PBS, placed into an enzymatic dissociation cocktail (2 U ml$^{-l}$ Liberase (Sigma-Aldrich, 5401127001) and 2 U ml$^{-1}$ elastase (Worthington, LS002279) in Hank's Balanced Salt Solution (HBSS) and minced. After incubation at 37°C for 1 h, the cell suspension was strained and then pelleted by centrifugation at 500g for 5 min. The enzyme solution was then discarded, and cells were resuspended in fresh HBSS. To increase biological replication, multiple mice were used to obtain single-cell suspensions at each time point. For each DRE-LacZ mice scRNA capture, 3 mice were used. Two replicates performed for control, and 1 replicate for TCDD treatment. For each SMC-LnT and SMC-AhrKO mice scRNA capture, 3 mice were used. Three replicates were performed for control, 2 replicates were performed for 16 weeks TCDD treatments, and 1 replicate for 8 weeks TCDD treatment. For each scATAC capture, 3 mice were used. Two replicates were performed for control, and 1 replicate for TCDD treatment. Cells were sorted FACS sorted based of tdTomato expression. tdT$^+$ cells (considered to be of SMC lineage) and tdT$^-$ cells were then captured on scRNA-Seq workflow and datasets were later combined for all subsequent analyses. For single cell ATAC, tdT$^+$ cells and tdT$^-$ cells were pooled at a 1:1 ratio, collected in BSA-coated tubes, and nuclei isolated per 10X recommended protocol, and captured on the 10X scATAC platform.

**Single-Cell capture, library preparation and sequencing**

All single-cell capture and library preparation was performed at the Stanford Functional Genomics Facility (SFGF). Cells were loaded into a 10X Genomics microfluidics chip and encapsulated with barcoded oligo-dT-containing gel beads using the 10X Genomics Chromium controller according to the manufacturer's instructions. Single-cell libraries were then constructed according to the manufacturer's instructions. Libraries from individual samples were multiplexed into one lane before sequencing on an Illumina platform.

**Analysis of scRNA data**

Fastq files from each experimental time point and mouse genotype were aligned to the reference genome individually using CellRanger Software (10X Genomics). Individual datasets were aggregated using the CellRanger aggr command without subsampling normalization. The aggregated dataset was then analyzed using the R package Seurat.[7] For DRE-LacZ mice, the dataset was trimmed of cells expressing fewer than 750 genes, and genes expressed in fewer than 3 cells. The number of genes, the number of unique molecular identifiers (UMIs) and the percentage of mitochondrial genes were examined to identify outliers. As an unusually high number of genes can result from a "doublet" event, in which two different cell types are captured together with the same barcoded bead, cells with < 3250 genes were discarded. Cells containing >30% mitochondrial genes were presumed to be of poor quality and were also discarded. For SMC-LnT and SMC-AhrKO mice, the dataset was trimmer of cells expressing fewer than 1500 genes and more than 6000 genes. Cells containing >10% mitochondrial genes were discarded. The gene expression values then underwent library size normalization, using the published *sctranform* function in the Seurat R package.[8] Principal component analysis was used for dimensionality reduction, followed by clustering in PCA space using a graph-based clustering approach. UMAP was then used for two-dimensional visualization of the resulting clusters. The "*ROS score*" was constructed using average expression of the following genes- *Sod1, Gpx1, Cat, Hmox2, Nfe2l1, Nfe2l2, Tnfa, Hif1a, Sirt1*. The "*M1 polarization*" gene set included *Nos2, Tnf, Il1b, Cd86, Cxcl9, H2-Ab1, Ccl2, Stat1*; and the "*M2 polarization*" gene set included *Arg1, Mrc1, Chi3l3, Fizz1, Il10, Ccl17, Cd163, Il4ra, Tgm2*. Raw data from single-cell RNAseq data that support the findings of this study will be deposited in the GEO database. Primary accession codes are pending. Wilcoxon rank sum test was used for identification of differential markers between clusters. Upstream TF analysis was performed using TRRUST, a manually curated database of transcriptional regulatory networks.[9]

**Analysis of scATAC data**
Analysis of scATAC was performed as previously described.[10] Briefly, fastq files from each experimental time point and mouse genotype were aligned to the reference ATAC genome (mm10) individually using CellRanger Software (10x Genomics). Individual datasets were aggregated using the CellRanger aggr command without subsampling normalization. The aggregated dataset was then analyzed using the R package Signac. The dataset was first trimmed of cells containing fewer than 1100 peaks. The subsequent cells were then again filtered based off TSS enrichment, nucleosome signal, and percent of reads that lies within peaks found within the larger dataset. Cells with greater than 70,000 reads within peaks or fewer than 1100 peaks, <3 TSS enrichment, or <30% reads within peaks were removed as they are likely poor-quality nuclei. The remaining cells were then processed using *RunTFIDF*, *RunSVD* functions from Signac to allow for latent semantic indexing (LSI) of the peaks, which was then used to create UMAPs. Differentially accessible peaks between different populations of cells were found using *FindMarker* function, using number of peaks as latent variable to correct for depth. Motif matrix was obtained from JASPAR 2020, aligned onto BSgenome.Mmusculus.UCSC.mm10. Accessibility analysis around different motif was performed using *ChromVar*. Merging of scRNA and ATAC data was performed using Pseudo-expression of each cell created using *GeneActivity* function to assign peaks to nearest genes expressed in the scRNA dataset and mapped onto each other using Canonical Correlation Analysis.

**Preparation of mouse aortic root sections**
Immediately after sacrifice, mice were perfused with 0.4% PFA. The mouse aortic root was excised and immersed in 4% PFA at 4℃ for 12 hours (for immunohistochemistry and immunofluorescence) to 24 hours (for RNAscope). After passing through a sucrose gradient, tissue was frozen in OCT to make blocks. Blocks were cut into 7um-thick sections for further analysis.

**Mouse aortic root histology**

Slides were prepared and processed as described previously.[11] For immunohistochemistry (IHC), sections were then incubated overnight at 4℃ with anti-SM22alpha (Tagln) rabbit polyclonal primary antibody (Abcam No. ab14106, 1:300 dilution), or CD68 rabbit polyclonal antibody (Abcam No. ab125212, 1:300 dilution). Sections were washed for 5 minutes x2 with TBS and then incubated with the Rabbit-on-Rodent HRP Polymer (Biocare Medical, RMR622) for 30 minutes at room temperature (RT). Sections were washed x2 with TBS and then incubated with the Betazoid DAB chromogen reagents (Biocare Medical, BDB2004) for 4 minutes at RT. Sections were washed x2 in DI water and air-dried, followed by mounting with Fluoroshiled with 4′,6-diamidino-2-phenylindole (DAPI) (Sigma, F6057). For C3 and CD11b (Itgam) immunofluorescence (IF), antigen retrieval with sodium citrate buffer was performed, and C3 antibody (Abcam No. ab11862) and CD11b (Itgam) antibody (Fisher Scientific, 557394) was used as primary antibody, Goat anti-mouse antibody (Thermo Fisher, A32723) was used as secondary antibody. For the alkaline phosphatase enzymatic assay, Ferangi Blue Chromogen kit (Biocare Medical, FB813H) was used as instructed. Researchers were blinded to the genotype of the animals until completion of the analysis. The IHC processed sections were visualized using Leica DM5500 microscope and images were obtained using Leica Application Suite X software. The IF processed sections were visualized using ECHO Revolve. Areas of interest were quantified using ImageJ (NIH) software and compared using a two-sided *t*-test. The lesion cap was defined as 30μm segment from the luminal surface as previously described.[11, 12]

**RNAscope**

Slides were processed according to the manufacturer's instructions, and all reagents were obtained from ACD Bio. Slides were washed once in PBS, then immersed in Target Retrieval reagent at 100 °C for 5 min. Slides were washed twice in deionized water, immersed in 100% ethanol and air dried, and sections were encircled with a liquid-blocking pen. Sections were incubated with Protease Plus reagent for 30 min at 40°C, then washed twice with deionized water. Sections were incubated with probes against mouse C3 or a negative control probe for 2 h at 40 °C. RNAscope HD assay (RED) and Multiplex colorimetric assay were performed per the manufacturer's instructions.

10.1016/j.molcel.2010.05.004. PubMed PMID: 20513432; PubMed Central PMCID: PMCPMC2898526.
6. Corces MR, Trevino AE, Hamilton EG, Greenside PG, Sinnott-Armstrong NA, Vesuna S, Satpathy AT, Rubin AJ, Montine KS, Wu B, Kathiria A, Cho SW, Mumbach MR, Carter AC, Kasowski M, Orloff LA, Risca VI, Kundaje A, Khavari PA, Montine TJ, Greenleaf WJ, Chang HY. An improved ATAC-seq protocol reduces background and enables interrogation of frozen tissues. Nat Methods. 2017;14(10):959-62. Epub 2017/08/29. doi: 10.1038/nmeth.4396. PubMed PMID: 28846090; PubMed Central PMCID: PMCPMC5623106.
7. Butler A, Hoffman P, Smibert P, Papalexi E, Satija R. Integrating single-cell transcriptomic data across different conditions, technologies, and species. Nat Biotechnol. 2018;36(5):411-20. Epub 20180402. doi: 10.1038/nbt.4096. PubMed PMID: 29608179; PubMed Central PMCID: PMCPMC6700744.
8. Hafemeister C, Satija R. Normalization and variance stabilization of single-cell RNA-seq data using regularized negative binomial regression. Genome Biol. 2019;20(1):296. Epub 20191223. doi: 10.1186/s13059-019-1874-1. PubMed PMID: 31870423; PubMed Central PMCID: PMCPMC6927181.
9. Han H, Cho JW, Lee S, Yun A, Kim H, Bae D, Yang S, Kim CY, Lee M, Kim E, Lee S, Kang B, Jeong D, Kim Y, Jeon HN, Jung H, Nam S, Chung M, Kim JH, Lee I. TRRUST v2: an expanded reference database of human and mouse transcriptional regulatory interactions. Nucleic acids research. 2018;46(D1):D380-D6. doi: 10.1093/nar/gkx1013. PubMed PMID: 29087512; PubMed Central PMCID: PMCPMC5753191.
10. Cheng P, Wirka RC, Shoa Clarke L, Zhao Q, Kundu R, Nguyen T, Nair S, Sharma D, Kim HJ, Shi H, Assimes T, Brian Kim J, Kundaje A, Quertermous T. ZEB2 Shapes the Epigenetic Landscape of Atherosclerosis. Circulation. 2022;145(6):469-85. Epub 20220106. doi: 10.1161/CIRCULATIONAHA.121.057789. PubMed PMID: 34990206; PubMed Central PMCID: PMCPMC8896308.
11. Wirka R, D. W, Paik DT, Pjanic M, Nguyen AT, Miller CJ, Kundu R, Nagao M, Coller J, Koyano TK, Fong R, Woo YJ, Liu B, Montgomery SB, Wu JC, Zhu K, Chang R, Alamprese M, Tallquist MD, Kim JB, Quertermous T. Single cell analysis of smooth muscle cell phenotypic modulation in vivo reveals a critical role for coronary disease gene TCF21 in mice and humans. Nat Med. 2019;25:1280-9.
12. Shankman LS, Gomez D, Cherepanova OA, Salmon M, Alencar GF, Haskins RM, Swiatlowska P, Newman AA, Greene ES, Straub AC, Isakson B, Randolph GJ, Owens GK. KLF4-dependent phenotypic modulation of smooth muscle cells has a key role in atherosclerotic plaque pathogenesis. Nat Med. 2015;21(6):628-37. doi: 10.1038/nm.3866. PubMed PMID: 25985364.
